# Ki-67 and condensins support the integrity of mitotic chromosomes through distinct mechanisms

**DOI:** 10.1101/202390

**Authors:** Masatoshi Takagi, Takao Ono, Toyoaki Natsume, Chiyomi Sakamoto, Mitsuyoshi Nakao, Noriko Saitoh, Masato T. Kanemaki, Tatsuya Hirano, Naoko Imamoto

**Affiliations:** Cellular Dynamics Laboratory, RIKEN; Chromosome Dynamics Laboratory, RIKEN; Division of Molecular Cell Engineering, NIG; IMEG, Kumamoto University; Department of Cancer Biology, The Cancer Institute of JFCR

**Keywords:** Ki-67, condensin, mitotic chromosome, AID

## Abstract

Although condensins play essential roles in mitotic chromosome assembly, Ki-67, a protein localizing to the periphery of mitotic chromosomes, had also been shown to make a contribution to the process. To examine their respective roles, we generated a set of HCT116-based cell lines expressing Ki-67 and/or condensin subunits that were fused with an auxin-inducible degron for their conditional degradation. Both the localization and the dynamic behavior of Ki-67 on mitotic chromosomes were not largely affected upon depletion of condensin subunits, and vice versa. When both Ki-67 and SMC2 (a core subunit of condensins) were depleted, ball-like chromosome clusters with no sign of discernible thread-like structures were observed. This severe defective phenotype was distinct from that observed in cells depleted of either Ki-67 or SMC2 alone. Our results show that Ki-67 and condensins, which localize to the external surface and the central axis of mitotic chromosomes, respectively, have independent yet cooperative functions in supporting the structural integrity of mitotic chromosomes.

**List of Abbreviations used:**

AID: auxin-inducible degron
DOX: doxycycline
FRAP: fluorescence recovery after photobleaching
IAA: indol-3-acetic acid
mACl: mAID-mClover
mACh: mAID-mCherry
NEBD: nuclear envelope breakdown
SMC: structural maintenance of chromosomes
STLC: S-Trityl-L-cysteine
topo IIα: topoisomerase IIα

## Introduction

During mitosis of animal cells, the nuclear envelope breaks down and chromatin surrounded by the nuclear envelope is now packaged into a discrete set of rod-shaped structures, known as mitotic chromosomes. This process enables different chromosomes to individualize, duplicated chromatids to resolve, and sister kinetochores to properly attach to the mitotic spindle, thereby ensuring the faithful segregation of genetic materials into daughter cells. Extensive studies during the past two decades have established that a class of multiprotein complexes, condensins, play central roles in mitotic chromosome assembly and segregation (Hirano, 2016; Uhlmann, 2016). Most eukaryote species have two different types of condensin complexes (condensins I and II). The two complexes share the same pair of structural maintenance of chromosome (SMC) ATPase subunits (SMC2 and SMC4), and have distinct sets of non-SMC regulatory proteins (CAP-H, -D2, and -G for condensin I, CAP-H2, -D3, and -G2 for condensin II). A recent study has shown that structures reminiscent of mitotic chromosomes can be reconstituted *in vitro* using a limited number of purified factors, including core histones, three histone chaperones, topoisomerase IIα (topo IIα), and condensin I (Shintomi et al., 2015). It is clear, however, that this list represents a minimum set of proteins required for building mitotic chromosomes, and that additional proteins must cooperate to provide them with physical and physicochemical properties that support and promote their own segregation. Candidates for such proteins include linker histones (Maresca et al., 2005), the chromokinesin KIF4 (Mazumdar et al., 2006; Samejima et al., 2012; Takahashi et al., 2016) and Ki-67 (Booth et al., 2016; Takagi et al., 2016).

Ki-67 is a nucleolar protein widely appreciated as a cell proliferation marker (Scholzen and Gerdes, 2000). During mitosis, Ki-67 is localized around mitotic chromosomes and constitutes a perichromosomal layer to which many nucleolar proteins are targeted (Booth et al., 2014; Takagi et al., 2014). To assess the mitotic function of Ki-67, we have recently generated HCT116-based cell lines in which endogenous Ki-67 was degraded conditionally and acutely via an auxin-inducible degron (AID) (Takagi et al., 2016). Using the cell lines, we demonstrated that Ki-67 aids the finalization of mitotic chromosome assembly and the maintenance of rod-shaped chromosome structures (Takagi et al., 2016). Another recent study has demonstrated that Ki-67 may act as a biological “surfactant” to prevent the coalescence of mitotic chromosomes by using its positively-charged, extended conformation that orients perpendicular to the surface of mitotic chromosomes (Cuylen et al., 2016). Despite these intriguing observations, it remains unclear how the perichromosomally localized proteins such as Ki-67 might functionally cooperate with the axially localized proteins such as condensins to build individual chromosomes and to support their segregation during mitosis.

In the current study, we aimed to address the question by conditionally depleting Ki-67 and condensin subunits individually or simultaneously from mitotic cells. To this end, we generated a panel of HCT116-based cell lines expressing Ki-67 and/or condensin subunits that were fused with AID for their conditional degradation and with fluorescent proteins for imaging. Remarkably, ball-like chromosome clusters with no sign of discernible thread-like structures were observed in mitotic cells depleted of both Ki-67 and SMC2. To further assess this unprecedented “slime-ball” phenotype, we introduced a quantitative analysis using a supervised machine-learning algorithm, wndchrm (Ono et al., 2017; Orlov et al., 2008). We also present evidence that abberant kinetochore-microtubule attachments accompany the formation of the slime ball. The observations presented here argue that Ki-67 and condensins, which localize to the external surface and the central axis of mitotic chromosomes, respectively, have independent yet cooperative functions in supporting the structural integrity of mitotic chromosomes in mammalian cells.

## Results

### Ki-67 and hCAP-H/H2 localize on mitotic chromosomes independently of one another

We previously generated an HCT116-based cell line (AID2) in which endogenous Ki-67 was fused to mAID and mClover (mACl), thereby enabling us to degrade Ki-67 conditionally upon addition of indol-3-acetic acid (IAA) (Takagi et al., 2016). To examine the localization of hCAP-H (a subunit specific to condensin I) and hCAP-H2 (a subunit specific to condensin II) in the absence of Ki-67, we further modified AID2, by a CRISPR-mediated knock-in strategy, to generate AID11 and AID44, in which hCAP-H and hCAP-H2, respectively, were C-terminally fused to mCherry. AID11 and AID44 cells were synchronized to G2 phase in the presence or absence of IAA, and then released into M phase in the presence of S-Trityl-L-cysteine (STLC), a KIF11/Eg5 inhibitor (Fig. 1A). One hour after the release, the cells were subjected to immunoblot analysis (Fig. 1B) and microscopic observation (Fig. 1C,D). In both cell lines, Ki-67 fused to mACl was degraded upon addition of IAA (Fig. 1B, lanes 3 and 6) as had been shown in their ancestor cell line AID2 (Takagi et al., 2016). Whereas hCAP-H-mCherry and hCAP-H2-mCherry were detected at positions bigger than their endogenous counterparts in the blots due to the mCherry-tagging, no signal was detected at the size of their endogenous counterparts (lanes 2-3 and 5-6), indicating that both alleles of their genomes had been edited as intended. In a subpopulation of IAA-treated cells (~10% at most), the fluorescence signal of Ki-67-mACl was still detectable under the current condition. In the microscopic observation, we focused on cells in which the signal decreased to an undetectable level. We found that both hCAP-H-mCherry and hCAP-H2-mCherry localized at the axial regions of mitotic chromosomes similarly to the endogenous counterparts even though the overall morphology of chromosomes slightly swelled upon Ki-67 degradation (Fig. 1C,D), a result consistent with the previous observation obtained by immunofluorescence using specific antibodies (Takagi et al., 2016).

**Figure 1.**
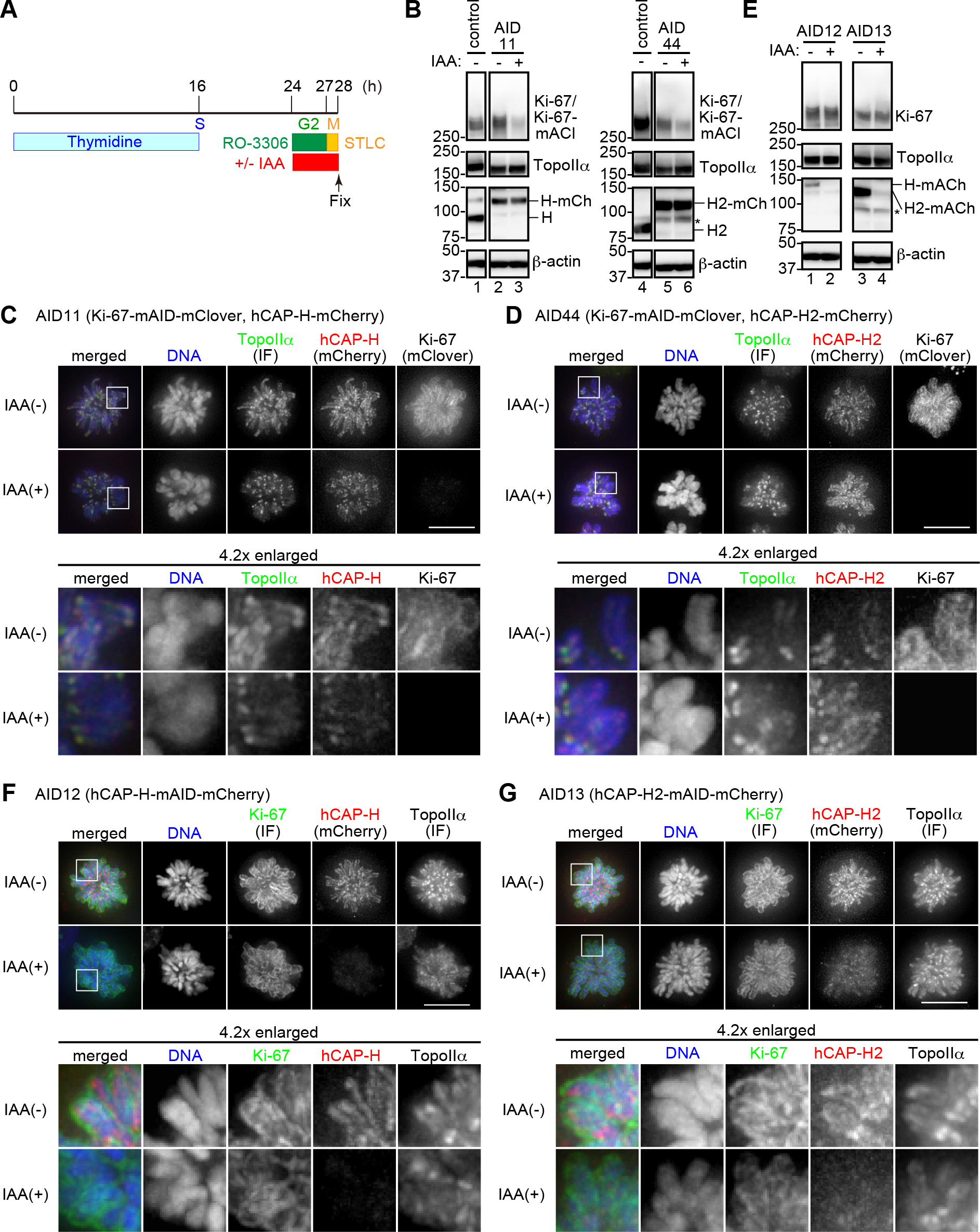
Ki-67 and hCAP-H/H2 localize on mitotic chromosomes independently of one another. (A) Schematic diagram of the cell preparation protocol. Thymidine (2 mM), RO-3306 (10 μM), STLC (10 μM), and IAA (0.5 mM) were added and/or washed out at the indicated time points. AID-tagged proteins are subjected to proteasome-mediated degradation upon the treatment of cells with IAA. (B) Immunoblot analysis of HCT116 (control), AID11 and AID44 cells. Membranes were probed with specific antibodies against the indicated proteins. The asterisk indicates a non-specific band. (C-D) Immunofluorescence analysis of AID11 and AID44 cells prepared in the absence (-) or presence (+) of IAA. Ki-67 and hCAP-H/H2 were detected via the fluorescence of mClover and mCherry, respectively, fused to their C-terminal ends. Topo IIα was detected by indirect immunofluorescence (IF) using a specific antibody. DNA was counterstained with Hoechst 33342. (E) Immunoblot analysis of AID12 and AID13 cells. Membranes were probed with specific antibodies against the indicated proteins. The asterisk indicates a non-specific band. (F-G) Immunofluorescence analysis of AID12 and AID13 cells prepared in the absence (-) or presence (+) of IAA. hCAP-H and hCAP-H2 were detected via the fluorescence of mCherry fused to their C-terminal ends. Ki-67 and topo IIα were detected by indirect immunofluorescence (IF) using specific antibodies. DNA was counterstained with Hoechst 33342. The areas indicated by the white squares are 4.2-times enlarged and shown on the bottom. Scale bars, 10 μm.

We then wished to investigate the localization of Ki-67 in the absence of hCAP-H or hCAP-H2. To this end, we generated AID12 and AID13 in which hCAP-H and hCAP-H2, respectively, were C-terminally fused to mAID and mCherry (mACh). Their conditional degradation upon addition of IAA was verified by immunoblot analysis (Fig. 1E) and microscopic observation (Fig. 1F,G). Upon depletion of either hCAP-H or hCAP-H2, the perichromosomal localization of Ki-67 was not significantly altered (Fig. 1F,G). Taken these results together, we conclude that Ki-67 and hCAP-H/H2 localize on mitotic chromosomes independently of one another.

### Ki-67 displays its perichromosomal localization and associates dynamically with mitotic chromosomes regardless of the presence or absence of SMC2

To determine unequivocally whether Ki-67 localizes to the periphery of mitotic chromosomes independently of condensins, we wish to generate a cell line in which SMC2, an ATPase subunit shared by condensins I and II, was C-terminally fused to mACh. We first attempted to generate such a cell line based on the standard protocol using NIG272 as a mother cell line, in which 0/TIR1, a ubiquitin E3 enzyme specific to AID-tagged proteins, was expressed constitutively, but without success, possibly due to reduced expression of SMC2 in the absence of IAA (discussed in Natsume et al. 2016). We circumvented this problem by using NIG430 as a mother cell line, in which ftsTIR1 was expressed conditionally upon addition of doxycycline (DOX) (Natsume et al., 2016). The resultant cell line, AID30, were treated according to the protocol shown in Fig. 2A, and then subjected to immunoblot analysis (Fig. 2B) and microscopic observation (Fig. 2C,D). We noticed that degradation of SMC2-mACh upon the treatment with IAA was inefficient in AID30, probably because of a lower expression level of 0/TIR1 compared to that seen in the mother cell (NIG430) (Fig. 2B). Reflecting this, variable levels of the SMC2-mACh signal were detected among individual cells treated with IAA (Fig. 2C,D). Cells where SMC2-mACh was reduced to less than 25% of the original level upon the treatment with IAA were rare (15 out of 127 cells: 11.8%; Fig. 2C). We noticed, however, that, in all cells with undetectable levels of SMC2-mACh, chromosomes lost their slim rod-like shapes and contracted into a smaller space (Fig. 2D, the third row). Although individualization of each chromosome became ambiguous in these cells, it was still possible to trace Ki-67 on the poorly organized chromosomes, indicating that SMC2 is largely dispensable for the perichromosomal localization of Ki-67.

**Figure 2.**
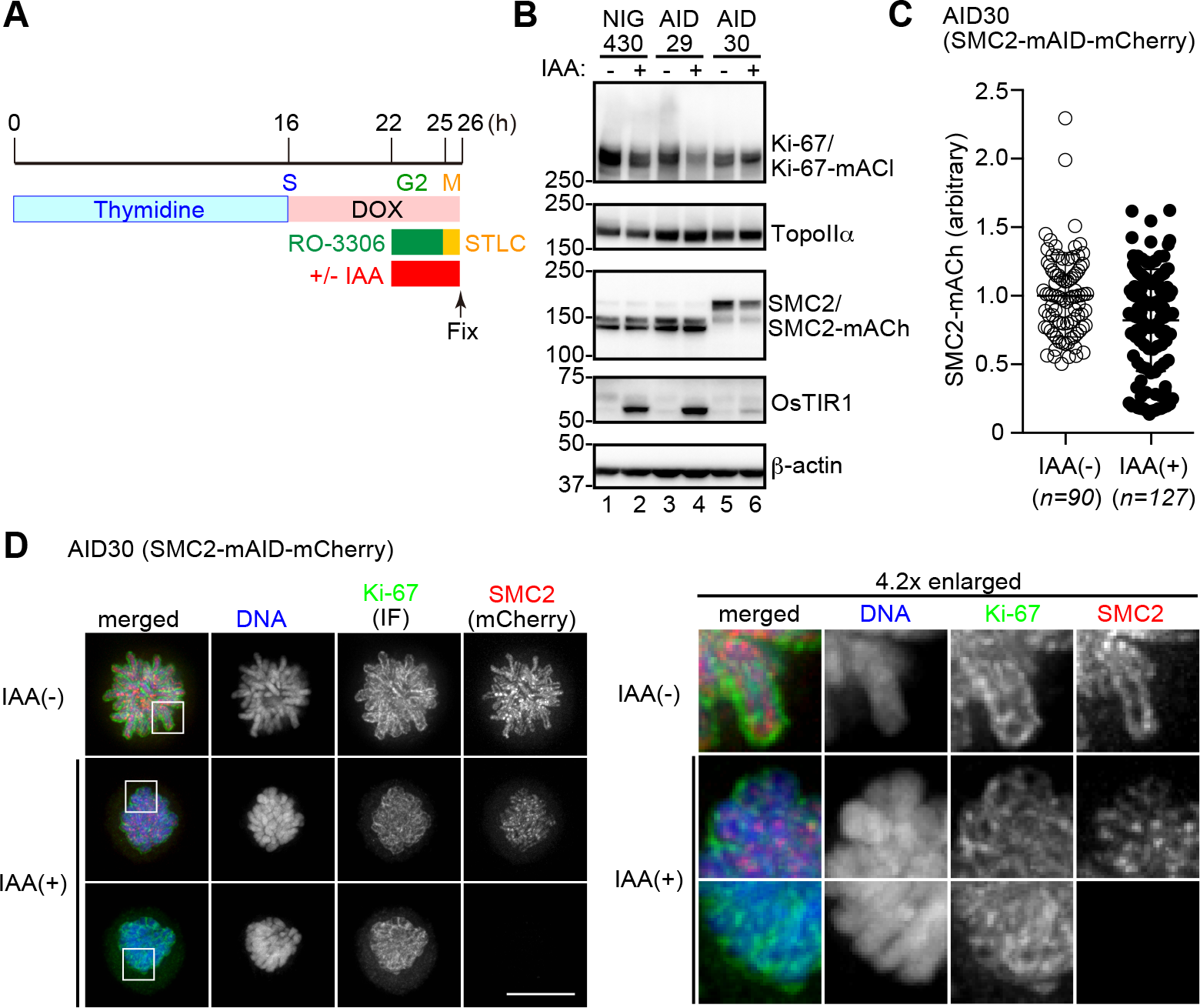
Ki-67 localizes on mitotic chromosomes independently of SMC2. (A)Schematic diagram of the cell preparation protocol. Thymidine (2 mM), DOX (2 μg/ml), RO-3306 (10 μM), STLC (10 μM), and IAA (0.5 mM) were added and/or washed out at the indicated time points. AID-tagged proteins are subjected to proteasome-mediated degradation upon the treatment of cells with DOX plus IAA. (B) Immunoblot analysis of NIG430 (a mother cell of AID29 and AID30), AID29 and AID30 cells. Membranes were probed with specific antibodies against the indicated proteins. (C) Fluorescence intensities of SMC2-mACh in AID30 cells prepared in the absence (-) or presence (+) of IAA. Each circle represents the fluorescence intensity of an individual cell relative to the average intensity of untreated AID30 cells. (D) Immunofluorescence analysis of AID30 cells prepared in the absence (-) or presence (+) of IAA. SMC2 was detected via the fluorescence of mCherry fused to the C-terminal end. Ki-67 and topo IIα were detected with indirect immunofluorescence (IF) using specific antibodies. DNA was counterstained with Hoechst 33342. Images of AID30 cells, in which SMC2-mAID-mCherry was degraded only moderately (second row) or completely (third row), are shown. The areas indicated by the white squares are 4.2-times enlarged and shown on the right. Scale bars, 10 μm.

We then wished to test whether depletion of SMC2 might affect this dynamic behavior of Ki-67 by fluorescence recovery after photobleaching (FRAP) experiments (Saiwaki et al., 2005). According to the protocol depicted in Fig. S2A, AID14 (expressing Ki-67-mACl plus hCAP-H-mACh; Fig. S1) cells were transfected twice with control siRNA (siControl) or siRNA against SMC2 (siSMC2), treated with reagents for synchronization, and then subjected to FRAP experiments. We measured the FRAP of Ki-67-mACl in cells with undetectable levels of hCAP-H-mACh (Fig. S2D,E), and found that it was indistinguishable from that observed in the control cells (Fig. S2B,C). These results indicate that the dynamic behavior of Ki-67 on the periphery of chromosomes does not depend on SMC2, or chromatin structure supported by condensins.

### Chromosomes rapidly lose their structural integrity upon nuclear envelope breakdown in cells devoid of both Ki-67 and SMC2

In the absence of either Ki-67 (Fig. 1C,D) or SMC2 (Fig. 2C), the architecture of mitotic chromosomes was compromised in different manners, suggesting that Ki-67 and condensins contribute to this event through distinct molecular mechanisms. To further examine the functional relationship between Ki-67 and condensins, we sought to deplete these chromosomal components simultaneously. To this end, we generated another cell line AID35, in which Ki-67 and SMC2 were C-terminally tagged with mACl and mACh, respectively. AID35 cells were treated according to the protocol shown in Fig. 3A. Cell lysates were prepared and subjected to immunoblot analysis to confirm that the bulk levels of the target proteins were substantially reduced in the presence of IAA (Fig. 3B). Microscopic inspection of individual cells revealed that the Ki-67-mACl levels were reduced to less than 20% of the original level in 59% of IAA-treated cells (19 out of 32 cells) inspected (Fig. 3C). Live cell imaging demonstrated that, in the absence of IAA, the behaviors of Ki-67-mACl and SMC2-mACh were indistinguishable from those of their endogenous counterparts in AID35 (Fig. 3D). We then monitored cells in which the signals of Ki-67 and SMC2 effectively disappeared in the presence of IAA (Fig. 3E). We found that chromosome compaction in the prophase nucleus was compromised in these cells (Fig. 3E, time −10’), as expected from the previous studies reporting condensin depletion (Ono et al., 2004; Hirota et al., 2004). A striking phenotype in chromosome morphology was observed immediately after nuclear envelope breakdown (NEBD). 10 min after NEBD and thereafter, chromatin formed a single cluster, displaying a ball-like structure in which the shape and border of individual chromosomes were not discerned (Fig. 3E, time 10’ −40’). To our knowledge, this type of abnormal chromosome structures in mitotic cells had not been reported in the literature before: we therefore refer to this unprecedented structure as a “slime ball” hereafter. Interestingly, the slime ball was frequently observed at one side of the cytoplasm, being placed at the vicinity of the cell cortex. Moreover, part of the slime ball seemed to be pulled in the opposite direction, forming “protrusions” that were readily discernible with Hoechst staining (see a later section).

**Figure 3.**
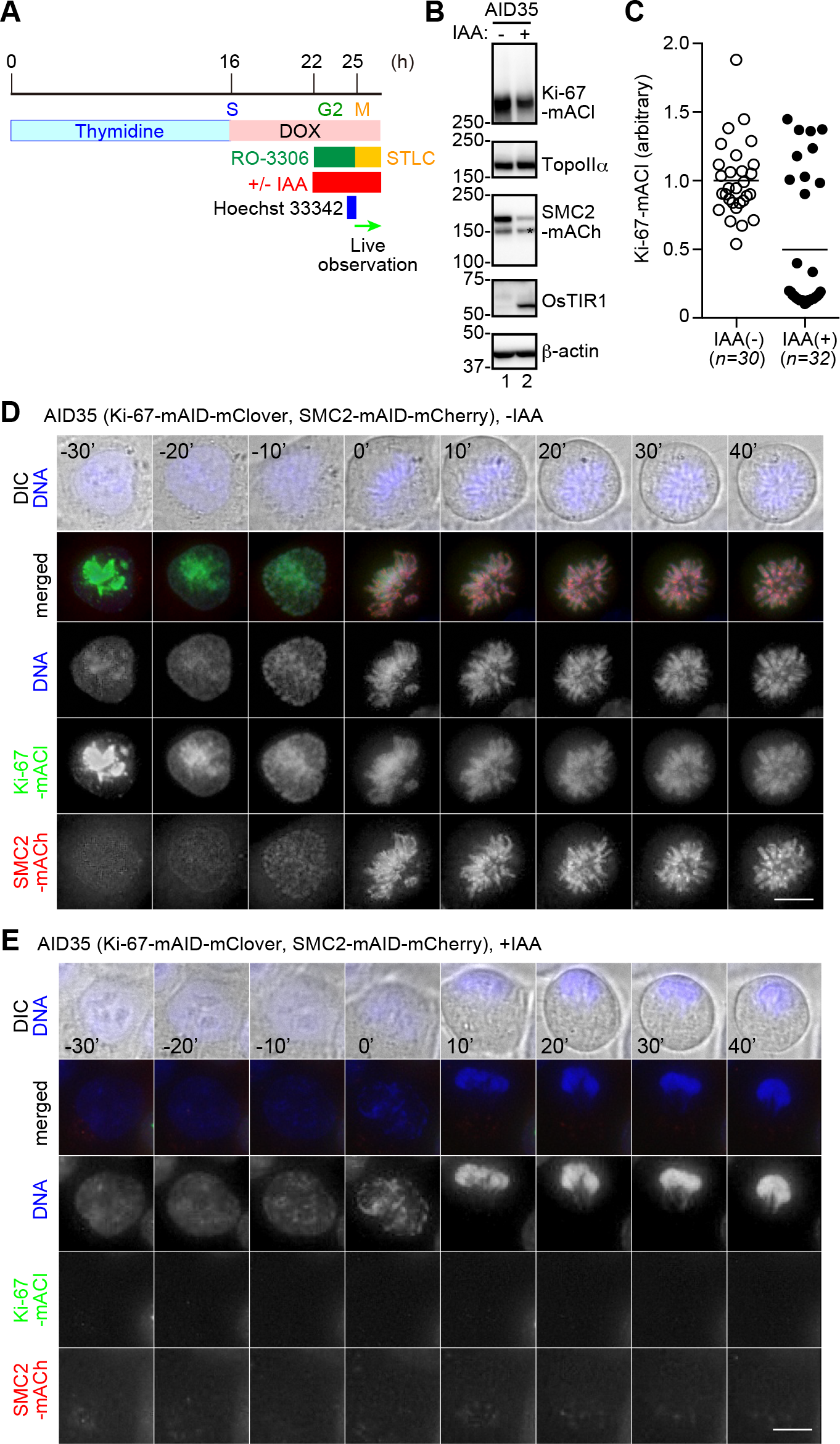
Rapid loss of the structural integrity of chromosomes immediately after NEBD in cells devoid of both Ki67 and SMC2. (A) Schematic diagram of the cell preparation protocol. Thymidine (2 mM), DOX (2 μg/ml), RO-3306 (10 μM), STLC (10 μM), Hoechst 33342 (100 ng/ml), and IAA (0.5 mM) were added and/or washed out at the indicated time points. (B) Immunoblot analysis of AID35 cells. Membranes were probed with specific antibodies against the indicated proteins. (C) Fluorescence intensities of Ki-67-mACl in AID35 cells prepared in the absence (-) or presence (+) of IAA. Each circle represents the fluorescence intensity of an individual cell relative to the average intensity of untreated AID35. The horizontal bars show the average intensities. (D-E) Live observation of AID35 cells prepared in the absence (D) or presence (E) of IAA. Images were taken at 10-min intervals. The first frame after NEBD marks the time point 0. DIC: differential interference contrast. Shown here are representative image sets out of more than six image sets captured. Scale bars, 10 μm.

### Mitotic chromosomes rapidly lose their structural integrity upon the degradation of both Ki-67 and SMC2 even after their assembly is complete

We next wished to test what would happen when the degradation of both Ki-67 and SMC2 was induced after mitotic chromosome assembly was complete. To this end, we treated AID35 cells with the protocol depicted in Fig. 4A. In this protocol, IAA was added one hour after removing RO-3306 (rather than being added at the same time of RO-3306 addition). At the time point when IAA was added, the cell had already entered mitosis, displaying a discrete set of condensed chromosomes: Ki67-mACl localized to the external surface of chromosomes, whereas SMC2-mACh was detectable on their central axis (Fig. 4B, time 0’). 40 min after addition of IAA, the signals of Ki-67-mACl and SMC2-mACh started to decrease, and chromosomes tended to lose their rod-shaped morphology (Fig. 4B, time 40’). After 60 min, both signals diminished to an undetectable level, and chromosomes form a single cluster with protrusions (Fig. 4B, time 60’, 80’ and 100’), whose morphology was very similar to that of the slime ball shown in Fig. 3. In control cells in which both Ki-67 and SMC2 escaped from degradation during the imaging period, mitotic chromosomes kept their structural integrity (Fig. S3C). Taken the observations in Figs. 3,4 together, we conclude that cells can neither establish nor maintain the structural integrity of mitotic chromosomes when both Ki-67 and SMC2 are absent.

**Figure 4.**
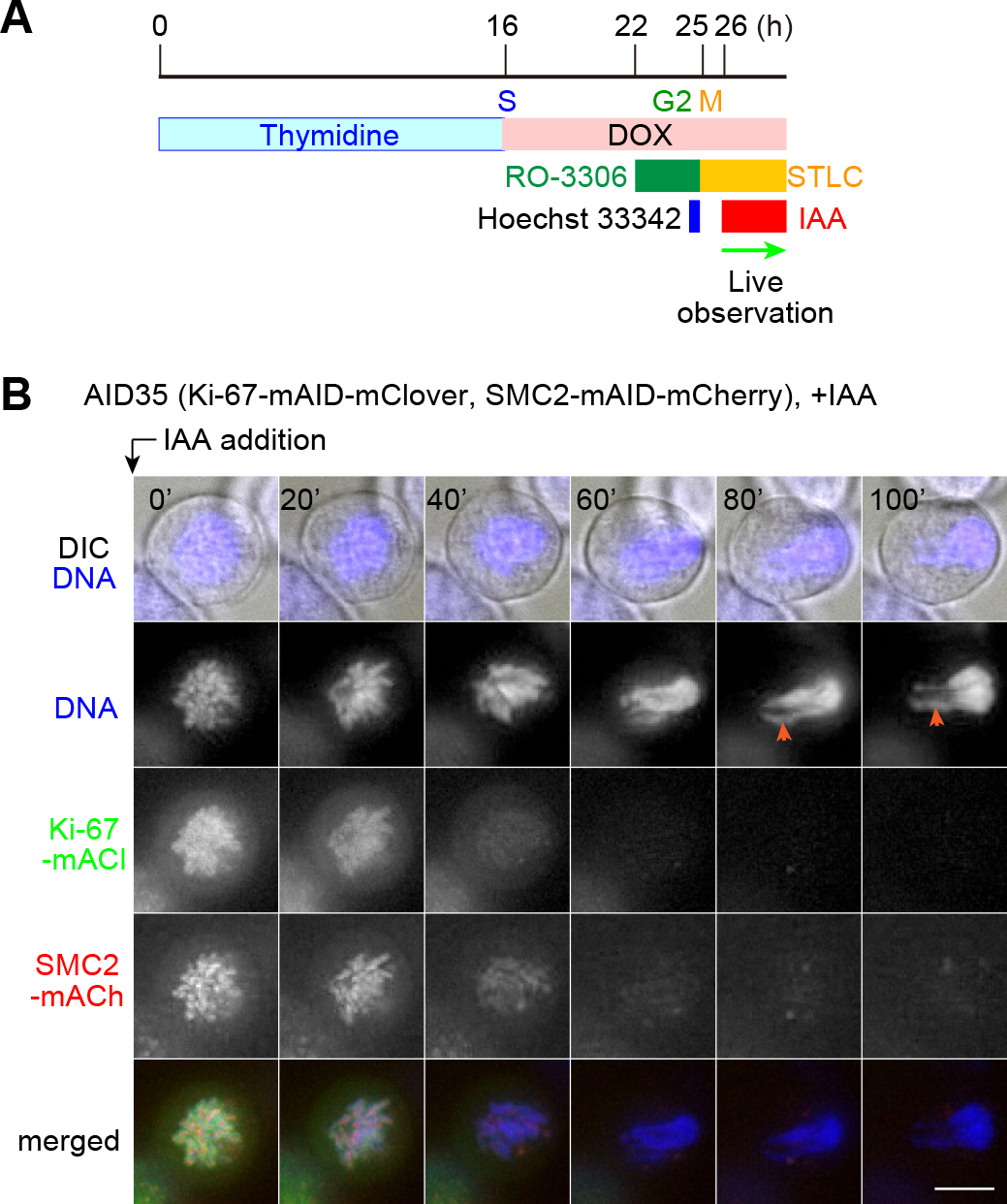
Rapid loss of the structural integrity of mitotic chromosomes in cells devoid of both Ki67 and SMC2 even after their assembly is complete. (A) Schematic diagram of the cell preparation protocol. Thymidine (2 mM), DOX (2 μg/ml), RO-3306 (10 μM), STLC (10 μM), Hoechst 33342 (100 ng/ml), and IAA (0.5 mM) were added and/or washed out at the indicated time points. AID-tagged proteins are subjected to proteasome-mediated degradation upon the treatment of cells with IAA. (B) Live observation of AID35 cells. Images were taken at 10-min intervals over 100 min, and only the selected frames are represented (all images are represented in Fig. S3). Ki-67-mACl and SMC2-mACh were degraded over time as intended, and mitotic chromosomes accordingly lost the structural integrity. The characteristic protrusions are marked with arrowheads. DIC: differential interference contrast. Shown here are representative image serieses of more than eight captured image serieses. Scale bars, 10 μm.

### Validation of the observed chromosome morphology by a machine learning algorism

In the experiments above, we observed seemingly different impacts on the morphology of mitotic chromosomes caused by depletion of either Ki-67, SMC2, or both of them. When comparing the images of those chromosomes side by side (Fig. 5A), the defective phenotypes observed among the three cell lines were clearly distinct from each other. Nevertheless, to further validate these differences objectively and quantitatively, we used a supervised machine-learning algorithm, wndchrm (weighted neighbor distances using a compound hierarchy of algorithms representing morphology) (Ono et al., 2017; Orlov et al., 2008). We first collected 36 images of mitotic chromosomes observed under each of the four conditions (control, Ki-67 depletion, SMC2 depletion, and Ki-67/SMC2 double depletion), and each set was defined as a class. Different numbers of images (5-35 images) were randomly selected from each class, and they were subjected to wndchrm analysis for constructing classifiers. We found, as expected, that the classification accuracy (CA) obtained with those classifiers increased according to the number of images used, reaching a plateau when more than 15 images from each class were used (Fig. 5B). Then, the 36 images in each class were randomly divided into two independent subclasses (subclasses 1 and 2, each containing 18 images). The resultant eight subclasses were processed in parallel for wndchrm analysis, and the differences included in those images were statistically evaluated. The results were displayed as morphological distances (MD) between two different subclasses (Fig. 5C) and also as a phylogeny tree (Fig. 5D). In the phylogeny tree, the two subclasses derived from each class were closely clustered with each other, confirming the assurance of this classification method. Notably, the four classes were distantly branched from each other, positioning at vertexes of a cruciform having four branches of similar lengths. This result indicates that the morphology of mitotic chromosomes formed in the four different settings tested are distinct from each other, and that the defective phenotype observed in cells devoid of both Ki-67 and SMC2 is closer to neither that observed in cells devoid of Ki-67 alone nor that observed in cells devoid of SMC2 alone.

**Figure 5.**
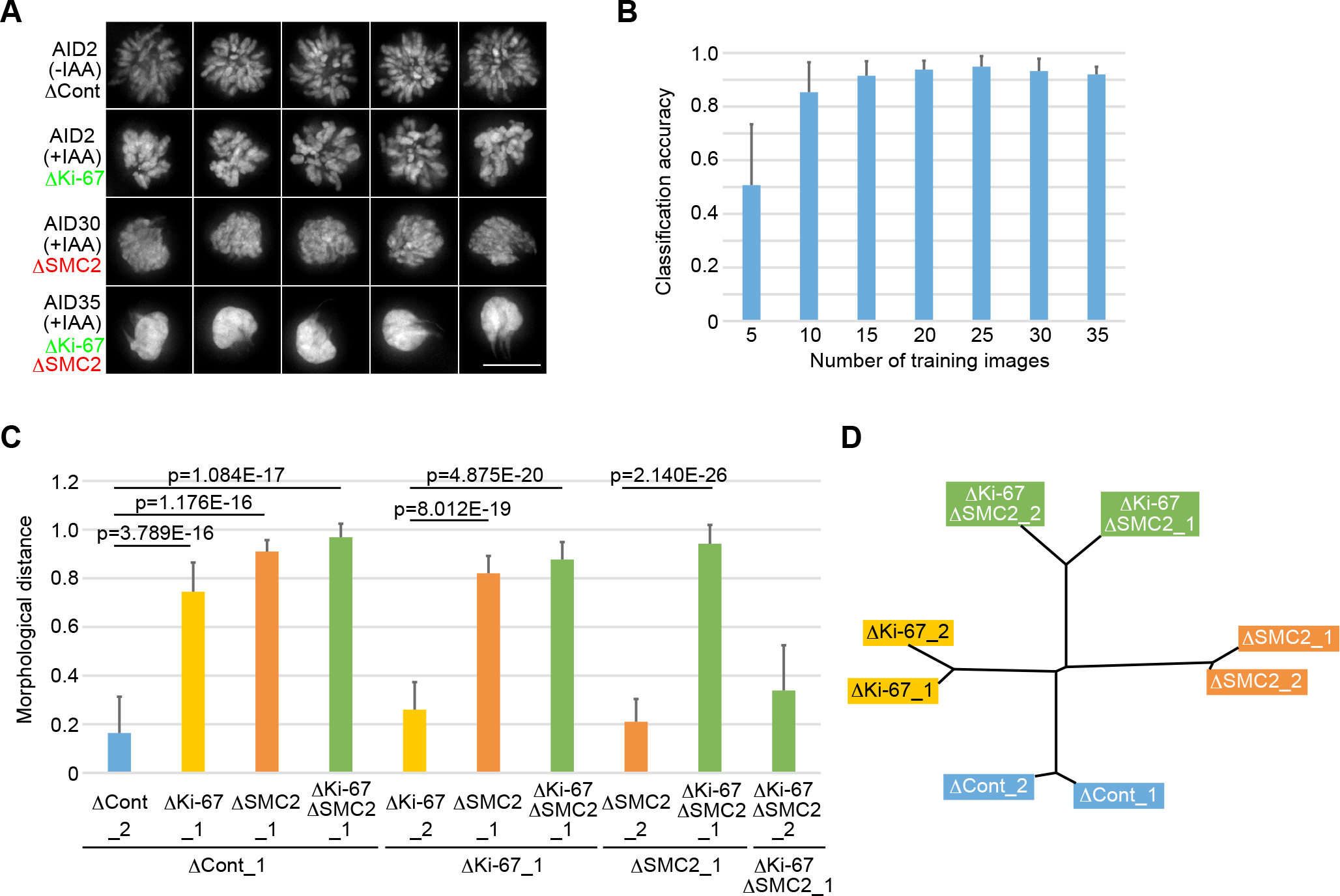
Quantitative analyses of chromosome morphology using a machine-learning algorism. (A) Representative images of mitotic chromosomes observed in four different experimental settings. The sources of the images include AID2 cells in the absence (-) or presence (+) of IAA according to the protocol illustrated in Figure 1 A, and AID30 and AID35 cells in the presence (+) of IAA according to the protocol illustrated in Figure 2 A. For collecting chromosome images from the IAA-treated cells, cells were selected in which the fluorescence signals of Ki-67-mACl and/or SMC2-mACh were diminished to an undetectable level. 36 images were collected from each setting and stored as four different classes (ACont, AKi-67, ASMC2 and AKi-67ASMC2). Scale bar, 10 αm. (B-D) Wndchrm analysis. (B) Assessment of the optimum numbers of training images required for classification. Different numbers of images (5-35 images) from each class were used as training images for constructing classifiers. The classification accuracy (CA) of each classifier was determined by cross validation tests. The values shown are the mean and SD from 20 independent tests. The accuracy reached a plateau when more than 15 training images were used. (C-D) Morphological distance (MD) and phylogeny. Images obtained from each of the four settings were randomly divided into two subclasses (each containing 18 images) and one of them served as a negative control of another. The MD values shown are the mean and SD from 20 independent cross validation tests. Statistics are from a two-tailed Student’s *t* test.

### Additional characterization of the slime-ball phenotype observed in cells devoid of both Ki-67 and SMC2

To further characterize the slime-ball phenotype observed in cells devoid of both Ki-67 and SMC2 (Fig. 3E), those cells were fixed one hour after the release into mitosis and subjected to immunofluorescence analyses using various antibodies (Fig. 6A-D). One of the most conspicuous observations upon depletion of both Ki-67 and SMC2 was the loss of radial arrays of microtubules and the emergence of microtubule bundles passing through the slime ball (Fig. 6A,D). The centrosomes, as judged by the localization of pericentrin, were located away from the chromosome mass (Fig. 6D). Chromosomal regions containing kinetochores, as determined by the localization of Hec1 (an outer kinetochore component), seemed to be pulled along the microtubule bundles toward the centrosome (Fig. 6B), thereby producing the protrusions characteristic of the slime ball. Interestingly, strong signals of topo IIα were detectable along the protrusions (Fig. 6C), implicating that topo II-enriched pericentromeric heterochromatic regions were also pulled by the microtubule bundles. In parallel with these observations, cells devoid of only SMC2 (AID30 treated with IAA) were examined (Figs. 6D,S4). Although Hec1 signals were seen clustered at one side of the nucleus as well (Fig. S4B) probably along microtubule fibers (Figs. 6D,S4A), accumulation of topo IIα on certain chromosomal regions was not clearly observed (Fig. S4C). The protrusions were less obvious in cells devoid of SMC2 alone compared to cells devoid of both Ki-67 and SMC2 (Figs. 6D,S4). In fact, the distance between the centroid of chromatin mass and the Hec1-positive region was shorter in the former cells than in the latter cells (Fig. 6E). Assuming that the emergence of the Hoechst-positive protrusions reflects a loss of structural integrity of chromosomes, the chromosomes devoid of both Ki-67 and SMC2 appeared more fragile than those devoid of SMC2 alone. The alteration of microtubule organization, which was accompanied with the depletion of SMC2, might contribute to and accelerate the formation of the slime-ball phenotype (see Discussion).

**Figure 6.**
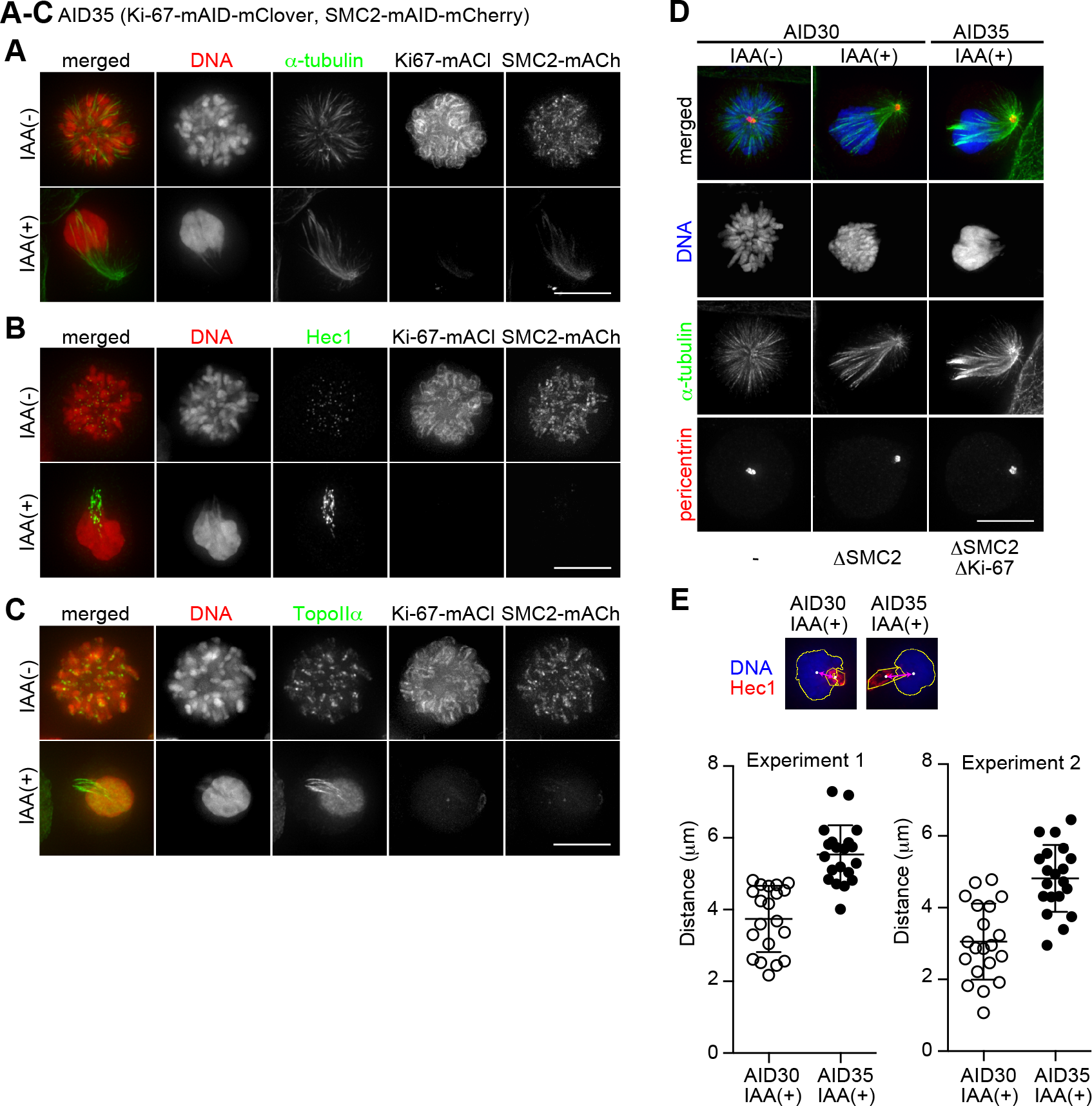
Behavior of chromosomal and non-chromosomal markers in cells devoid of both Ki-67 and SMC2. AID35 cells (A-C) and AID30 cells (D) were prepared in the absence (-) or presence (+) of IAA according to the protocol depicted in Figure 2 A, and processed for immunofluorescence using antibodies against α-tubulin (A), Hec1 (B), topo IIα (C), or α-tubulin and pericentrin (D). (A-C) Ki-67 and SMC2 were detected via the fluorescence of mClover and mCherry, respectively, fused to their C-terminal ends. (A-D) DNA was counterstained with Hoechst 33342. Scale bars, 10 αm. (E) Distances between the centroid of chromatin mass (blue) and Hec1-positive region (red) of IAA-treated AID30 and AID35 cells (20 cells each) were measured and plotted. The results from two independent experiments are shown.

**Figure 7.**
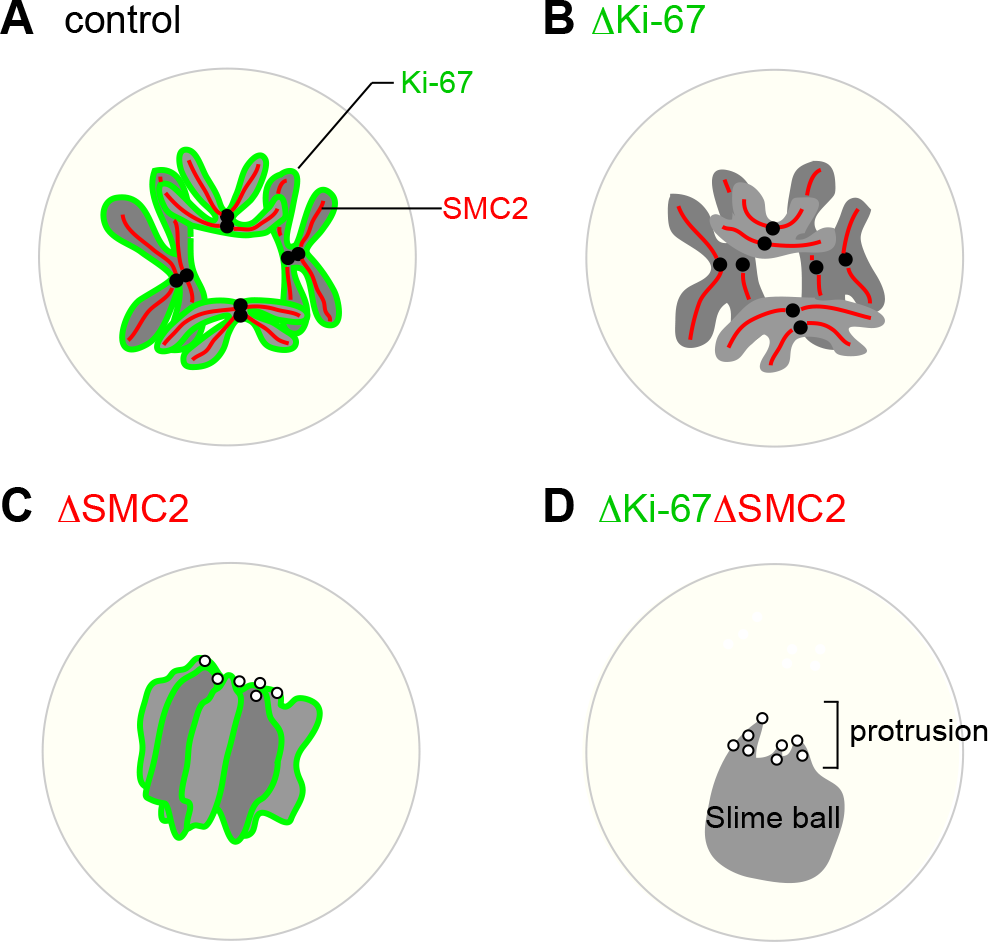
Ki-67 and condensins support the integrity of mitotic chromosomes through distinct mechanisms. (A) In control cells, Ki-67 (green) and condensins (red) support the integrity of mitotic chromosomes internally and externally, respectively. (B) In cells devoid of Ki-67, mitotic chromosomes are slightly swollen and tend to be coalesced with each other. (C) In cells devoid of SMC2, mitotic chromosomes are severely disorganized. The centromere/kinetochore regions tend to be clustered at the side of the chromatin mass close to the centrosome (not shown). (D) When both Ki-67 and SMC2 are depleted, all chromosomes apparently fused to form a single cluster, which we call a “slime ball”, from which the centromere/kinetochore regions tend to
protrude toward a single direction through the action of microtubules. See the text for details.

## Discussion

In the current study, we aimed to understand the functional relationship between Ki-67 and condensins in establishing and maintaining the structural integrity of mitotic chromosomes in human cells. Using a panel of HCT116-based AID cells, in which Ki-67 or subunits of condensins can be degraded conditionally, we first extended our previous finding that Ki-67 and condensins behaved independently in mitotic cells (Takagi et al., 2016). We then examined defective phenotypes caused by depletion of both Ki-67 and condensins. The defect we observed was unprecedentedly drastic, which was further validated by the image analyses using a supervised machine-learning algorithm, wndchrm. Our results suggest that Ki-67 and condensins have independent yet cooperative functions in supporting the structural integrity of mitotic chromosomes.

### Contribiution of Ki-67 to the structure of mitotic chromosomes is rather cryptic

In the current study, we have shown that mitotic chromosomes assembled in the absence of Ki-67 display a swollen morphology (Figs. 1,5; Takagi et al., 2016). Consistently, a recent study showed that the total volume of mitotic chromosomes (DAPI-stained chromatin regions) increased by 38% upon siRNA-mediated depletion of Ki-67 in RPE-1 cells (Booth et al., 2016). Somewhat inconsistent with these data, however, we and others also noticed that Ki-67 depletion had little impact on the morphology of mitotic chromosomes when they were spread on the surface of slide glasses (“mitotic spreads”) or when they were simply exposed to hypotonic buffers (Cuylen et al., 2016; data not shown). We infer that Ki-67’s contribution to the structural integrity of mitotic chromosomes becomes apparent only when they are in close proximity in the cell, and that this situation is heavily perturbed when the cells are exposed to hypotonic buffers and/or subjected to spreading techniques. This idea is consistent with the recent proposal that Ki-67 could function as a “surfactant” (electrostatic charge barrier) that helps to prevent the coalescence of individual chromosomes (Cuylen et al., 2016).

It deserves to mention that the defective impact on chromosome appearance upon depletion of Ki-67 was seen more evidently in the experiments using the AID system (Takagi et al., 2016; this study) than in the previous experiments using siRNAs (Takagi et al., 2014; Vanneste et al., 2009). This might be explained by the quickness of Ki-67 degradation which was realized by the AID system, and also by the sure selection of cells to be observed based on the loss of fluorescence (derived from the fluorescent protein tagged tandemly to the AID). Likewise, the AID system is also powerful in perturbing condensin functions (Figs. 2,5,S4): depletion of condensins with other methods used so far, such as siRNA transfection or transcriptional repression, has been reported to cause relatively mild defects in chromosome condensation (Gassmann et al., 2004).

### Double depletion of Ki-67 and condensins causes unprecedented severe defects in chromosome architecture and behaviors

The current study has shown that cells devoid of both Ki-67 and SMC2 fail to assemble thread-like mitotic chromosomes, instead forming a ball-like chromatin cluster with no discernible borders between individual chromosomes, which we call the “slime ball” (Fig. 3D). This particular phenotype was different from, and much severer than, that observed in the absence of either Ki-67 or SMC2 alone (Fig. 5A). We have also shown that the depletion of Ki-67 and SMC2 “after” the completion of chromosome assembly causes a drastic contration of thread-like chromosomes into a chromatin cluster reminiscent of the slime ball (Figs. 4,S3). Such a drastic phenotype has never been observed by depleting either Ki-67 (Takagi et al., 2016) or SMC2 alone after the completion of chromosome assembly. Together also with their independent behaviors in mitotic cells (Figs. 1,2,S2), it is reasonable to speculate that Ki-67 and SMC2 support the structural integrity of mitotic chromosomes through distinct molecular mechanisms. What is the relationship between the two distinct mechanisms? The subunits of condensins were widely conserved among eukaryotes and play fundamental roles in the organization and segregation of mitotic chromosomes (Hirano, 2016). On the other hand, the orthologs of Ki-67 are detectable only in vertebrates. Ki-67 could have evolved to play an auxiliary role in increasing the fidelity of segregation of chromosomes, especially, of large size. Such an evolutionary situation could parallel the emergence of increasing numbers of phase-separeted organelles in complex organisms (Banani et al., 2017). Alternatively, non-vertebrate cells could have a peripheral chromosomal protein that plays an equivalent role to that of Ki-67 but has no sequence similarity to it.

Vertebrate cells have two different condensin complexes, condensins I and II, which behave and function differently (Green et al., 2012; Hirota et al., 2004; Ono et al., 2004). To get more insight into the mechanism behind the formation of the slime-ball phenotype, we co-depleted Ki-67 with either hCAP-H or hCAP-H2 (Fig. S1). Double depletion of Ki-67/hCAP-H or Ki-67/hCAP-H2 displayed less severe defects than double depletion of Ki-67/SMC2: the chromosomes were more swollen than Ki-67-depleted chromosomes, but never produced a phenotype reminiscent of the slime-ball phenotype observed in cells devoid of both Ki-67 and SMC2. Thus, loss of both functions of condensins I and II, along with loss of Ki-67, is required to create the highly characteristic defective phenotype.

### What mechanism might underlie the formation of the slime ball?

In the current study, we have shown that depletion of SMC2 alone or Ki-67 and SMC2 commonly causes microtubules to lose their radial arrays and to make bundles in STLC-treated mitotic cells (Figs. 6A,D, and S4A). This defect in microtubule orientation is therefore specific to loss of SMC2, but not that of Ki-67. A similarly characteristic microtubule orientation was observed in STLC-treated cells when “end-on” attachments of microtubules to kinetochores were blocked by depleting Nuf2 (Silk et al., 2009), suggesting strongly that kinetochores formed in cells devoid of SMC2 are “laterally” attached to microtubule fibers. Consistent with the notion, α-tubulin and Hec1 showed localization patterns close to but exclusive from each other in cells devoid of SMC2 (Fig. S4D). It should be noted that the slime-ball phenotype was observed only in cells devoid of both Ki-67 and SMC2. While the slime ball as a whole was pushed to one side in the cytoplasm to the vicinity of the cell cortex, the kinetochores were clustered at the opposite side (Figs. 6B,S5) and pulled to the direction to the minus end of microtubules (Fig. 6D) probably via their lateral attachment to the bundled microtubules (Fig. S5D). Although the similar distribution of kinetochores relative to microtubule bundles occurred also in cells devoid of SMC2 alone (Fig. S4), those kinetochores tended to stay close to the chromatin mass (Fig. 6E). We imagine that the formation of the slime ball might be based on the reduced rigidity of chromosomes, which is caused by double depletion of Ki-67 and SMC2, and accelerated by the “unidirectional” pulling force exerted on kinetochores along the bundled microtubules, which is caused by depletion of SMC2 alone, as illustrated in Fig. S5E.

We verified that not only Hec1 (an outer kinetochore component; Fig. 6B) but also CENP-A (a centromeric chromatin component), CENP-I/hMis6 (an inner kinetochore component) and BubR1 (a spindle check point kinase) were localized in the protrusions of slime ball (Fig. S5). These obsrevations suggest that most, if not all, components of kinetochores remain intact within each kinetochore unit. We speculate, however, that some specific components or abilities necessary for the end-on attachments of microtubules are lost from the kinetochores in the slime ball (and similarly from those in cells devoid of SMC2 alone). It will be important to understand in the future how loss of SMC2 produces such a drastic and specific phenotype in kinetochore-microtuble attachments.

## Conclusions

The key observations we made in the current study are summarized in Fig. 7. The most important finding is that cells devoid of both Ki-67 and condensins rapidly lose the structural integrity of mitotic chromosomes to an unprecedented level. In light of this finding, we propose a new concept in which Ki-67 and condensins, which localize to the external surface and the central axis of mitotic chromosomes, respectively, cooperate to support the structural integrity of mitotic chromosomes through distinct mechanisms. Additionally, we propose the possibility that the contacts and interferences among mitotic chromosomes must be restricted through the action of Ki-67, otherwise the chromosome morphology is adversely affected.

## Materials and methods

### Establishment of cell lines and their handling

HCT116 cells and its derivatives were cultured at 37°l of cells, collectively called AID cells in the current manuscript, were generated from HCT116 cells via successive uses of CRISPR/Cas9-mediated genome editing as described previously (Natsume et al., 2016). Briefly, as the first step, the constitutive or DOX-inducible expression units of 0/TIR1 was integrated in the AAVS1 locus of the HCT116 genome to generate cells called NIG272 or NIG430, respectively (Natsume et al., 2016). In these cells, as the second step, cassette sequences encoding variable tags were knocked-in immediately upstream of the stop codons of genes to be analyzed. Table S1 contains a list of the parental lines, the target genes, the kind of tags, the sequences of guide RNAs, the plasmid names of targeting and knock-in constructs, and antibiotics used for generating AID cells. Of cellular clones selected by their resistant to antibiotics (700 μg/ml neomycin or 100 μg/ml hygromycin B), the final selection of AID11 and AID44 were carried out visually with fluorescence microscopy. For other cell lines, clones in which the genome had been edited as designed in both alleles were selected by genomic PCRs using KOD-plus-Neo (TOYOBO, Osaka, Japan) and appropriate primer sets listed in Table S2. Successful editing in both alleles was further confirmed by immunoblotting (to check the loss of target proteins of their original size).

For fixed-cell immunofluorescence, cells were seeded on coverslips treated with 10 μg/ml fibronectin (Wako, Osaka, Japan). For live-cell imaging, including FRAP analysis, cells were seeded onto glass-bottom dishes (IWAKI, Tokyo, Japan). 2 × 10^5^ cells were plated on a 35-mm dish (either a glass-bottom dish or a polystyrene dish containing four fibronectin-coated round coverslips) 1 day before the experiments and processed as follows. Cells were treated with 2 mM thymidine for 16 h, released in thymidine-free medium for 6-7 h, treated with 10 μM R0-3306 (Tocris, Minneapolis, MN) for 3 h for arrest at the G2/M boundary, and released in medium containing 10 μM STLC (Tokyo Chemical Industry, Tokyo, Japan) for arrest in mitosis. Cells were mock-treated or treated with 0.5 mM IAA (Tokyo Chemical Industry) during the period depicted in Figs. 1A,2A,3A,4A,S3A. For the observations of AID cells derived from NIG430 (AID29, AID30 and AID35), incubation with 2 μg/ml DOX (MP Biomedicals, Santa Ana, CA) after the release from thymidine block was added for inducing the expression of *Os*TIR1. For probing DNA in live-cell observations, cells were treated with 100 ng/ml Hoechst 33342, 30 min before removing RO-3306.

### Immunoblotting

Cells were washed twice with ice-cold PBS supplemented with 0.3 mM PMSF, collected by centrifugation, and snap-frozen in liquid nitrogen. Cell pellets were resuspended in buffer B (20 mM Tris-HCl [pH 7.5], 150 mM NaCl, 5 mM MgCl_2_, 0.1% NP-40, 1 mM DTT, Complete Protease Inhibitor Mixture [Roche, Basel, Switzerland], and PhosSTOP [Roche]) supplemented with 0.25 units/ml Benzonase (Novagen, Madison, WI, USA), kept on ice for 30 min, mixed with the same volume of 4x concentrated sample buffer (250 mM Tris-HCl [pH 6.8], 8% SDS, 40% glycerol, 0.02% bromophenol blue, and 0.1 M DTT), and heated at 95°C. The denatured protein samples were electrophoretically separated on a SuperSep Ace 5-20% gradient gel (Wako, Osaka, Japan) and blotted onto a Immobilon-P membrane (Merck Millipore, Billerica, MA). The following antibodies were used as primary antibodies at the indicated dilutions or concentrations: mouse anti-“-actin (1:5,000, AC-15; Sigma-Aldrich, St. Louis, MO), rabbit anti-NCAPH/hCAP-H (1:1,000, 11515-1; ProteinTech, Rosemont, IL), rabbit anti-hCAP-H2 (1 [ig/ml, AfR205-4L; Ono et al., 2003), rabbit anti-Ki-67 (1:1,000, sc-15402; Santa Cruz, Dallas, TX), rabbit-ftsTIR1 (1:1,000; Natsume et al., 2016), rabbit anti-SMC2 (1:1,000, ab10412; Abcam, Cambridge, UK), and mouse anti-topoisomerase II< (1:2,000, 1C5; MBL, Nagoya, Japan) antibodies. The following antibodies were used as secondary antibodies at the indicated dilutions: goat anti-mouse HRP (1:3,000, 170-6516; Bio-Rad, Hercules, CA), and goat anti-rabbit HRP (1:3,000, 170-6515; Bio-Rad) antibodies. Protein bands were visualized by chemiluminescence using Immobilon Western (Merck Millipore).

### Immunofluorescence

One hour after the removal of RO-3306, cells were fixed with 3.7% PFA in PBS at room temperature for 10 min. The fixed cells were permeabilized with 0.5% Triton X-100 in PBS for 5 min, blocked with a blocking solution (PBS containing 5 mg/ml BSA and 50 mM glycine) for 1 h, and processed for immunofluorescence. The following antibodies were used as primary antibodies at the indicated dilutions: mouse anti-μ-tubulin (1:10,000, DM1A; Sigma-Aldrich), mouse anti-BubR1 (1:400, 8G1; MBL), mouse anti-CENP-A (1:200, 3-19; MBL), rat anti-CENP-I/hMis6 (1:100, PD032; MBL), mouse anti-HEC1 (1:1,000, 9G3; GeneTex, Irvine, CA), rabbit anti-Ki-67 (1:200, sc-15402; Santa Cruz), mouse anti-Ki67 (1:500, NA-59; Merck Millipore), rabbit anti-pericentrin (1:1,000, ab4448; Abcam), and mouse anti-topoisomerase IIμ (1:1,000, 1C5; MBL) antibodies. Secondary antibodies conjugated with Alexa Fluor 488/594/647 were purchased from Thermo Fisher Science (Waltham, MA). DNA was counterstained with 0.5 μg/ml Hoechst 33342. Immunofluorescence images were captured with a DeltaVision Core (Applied Precision, Issaquah, WA, US) with an inverted microscope (IX71; Olympus, Tokyo, Japan), an UPlanApo 60×/1.40 objective lens (Olympus), and a CoolSNAP HQ2 camera (Photometrics, Tucson, AZ, US). Images from z sections spaced 0.5-μm apart were acquired, deconvolved with softWorx (Applied Precision), and presented as maximum intensity projections.

### Quantification of fluorescence intensities

Cells were fixed and stained with Hoechst 33342 as described above. Images of chromatin stained with Hoechst 33342 and the fluorescent proteins to be quantified were captured and processed as described above except for the use of an UPlanFL40×/0.75 objective lens (Olympus). Chromosomal regions were determined based on the Hoechst-stained images, and the total pixel intensities of fluorescence images from those regions were calculated. The obtained values were normalized by the average value of control cells (cells untreated with IAA) and plotted using GraphPad Prism6 (GraphPad Software, La Jolla, CA, US).

### FRAP

For FRAP experiments, we first attempted to generate a cell line expressing Ki-67-mClover plus SMC2-mACh, but without success. We therefore decided to use AID14 (a cell line expressing Ki-67-mACl plus hCAP-H-mACh) which had been generated for other purposes (Fig. S1). It has to be mentioned that SMC2 is not tagged with mAID in AID14. FRAP experiments were performed on the laser scanning confocal microscope FV1200 (Olympus) equipped with PLAPON 60XO/1.42 (Olympus). Cells expressing Ki-67-mACl and hCAP-H-mACh from their intrinsic promotors (AID14) were transfected with siControl (5’-CGUACGCGGAAUACUUCGAdTdT; Elbashir et al., 2001) or siSMC2 (5’-UGCUAUCACUGGCUUAAUdTdT; (Gerlich et al., 2006)) at 0 and 24 h at a final concentration of 10 nM using Lipofectamine RNAiMAX (Thermo Fisher Scientific), and synchronized in mitosis as described above. The cells were transferred to a humidified environmental chamber (Stage Top Incubator; TOKAI HIT, Shizuoka, Japan) maintaining its temperature at 37°C and the CO_2_ concentration at 5%, and subjected to FRAP analysis within 100 min after the release from the cell cycle arrest with RO-3306. One pre-bleach frame followed by 2-sec bleach time with 473 nm laser line at 80% transmission, and 8-10 post-bleach frames were recorded at 30-sec intervals. In parallel with the signal of Ki-67-mACl, the signals of hCAP-H-mACh and DNA (stained with Hoechst 33342) were recorded. The mean mClover fluorescence intensities of the bleached chromatin region for each time point was normalized to that of unbleached chromatin region at the same time point within the same cell. As cells tended to move slightly during the imaging time, the measurement areas were corrected manually relying on the chromatin images. The value for each time point was further normalized with that at the pre-bleached frame.

### Live cell observations

Cells cultured in a glass-bottom dish were mounted on an inverted microscope (IX71, Olympus) equipped with a humidified environment chamber (MI-IBC, Olympus) to maintain its temperature at 37°C and the CO_2_ concentration at 5%. Fluorescence images were collected with a DeltaVision Core (Applied Precision) from z sections (5 sections spanning 8 μm for Fig. 3; 5 sections spanning 2 μm for Figs. 4 and S3) every 10 min with 2 × 2 binning and presented as maximum intensity projections. Differential interference microscope images were acquired in parallel from a single focal plane.

### Morphological quantification of chromosome with wndchrm

Microscopic images of chromosomes were obtained from fixed cells stained with Hoechst 33342 using DeltaVision Core (Applied Precision) with an UPlanApo 60×/1.40 objective lens (Olympus). Images from z sections (24 sections spanning 11.5 μm) were obtained, deconvolved, and presented as maximum intensity projections. For quantitative assessment of chromosome structures, a supervised machine-learning algorithm, wndchrm (weight neighbor distance using a computed hierarchy of algorithms representing morphology) ver. 1.52 (Ono et al., 2017; Orlov et al., 2008; Tokunaga et al., 2014), was applied to 36 projected images (188×188 pixels, 8 bit) in each condition, which was defined as a class. All of the images in the defined class were applied to wndchrm, and morphological feature values were assigned by training a machine. Phylogenies were computed using the Fitch-Margoliash method implemented in the PHYLIP package ver.3.696, which was based on pairwise class similarity values reported by wndchrm ver. 1.52 (Felsensein, 1989; (Johnston et al., 2008)). For each analysis, cross-validation tests were automatically repeated for 20 times with 13 training/5 test image data set. The options used for the image analysis were a large feature set of 2919 (-l) and multi-processors (-m). To measure pairwise class dissimilarity, morphological distances (MD) were calculated as the Euclidean distances 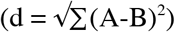 from the values in class probability matrix obtained from the cross-validations (Johnston et al., 2008). To calculate P values, two-sided Student’s *t*-test was performed for each of comparisons. To optimize the classification capacity, we measured classification accuracy (CA) using different numbers of training data sets, and found that the accuracy reached a plateau with more than 15 images (Fig. 5B). Then, each image in a class (36 images) was randomly assigned to two independent sets (folder 1 and folder 2, each containing 18 images) to confirm that images within the same class (condition) show negligible differences. They were expected to localize closely in phylogenies and to show low MD between them.

### Measurement of the distance between chromatin mass and Hec1-distributed region

Microscopic images of chromosomes and Hec1were obtained using DeltaVision Core (Applied Precision) with an UPlanApo 60×/1.40 objective lens (Olympus). Images from z sections (40 sections spanning 7.8 μm) were obtained, deconvolved, and presented as maximum intensity projections under the same condition. Chromosomal regions were determined by thresholding the chromosomal images. To delineate Hec1-positive regions, the convex hull of Hec1 signals were determined manually from the uniformly binarized images of Hec1. The distances between the centroids of these two regions were measured and plotted using GraphPad Prism6 (GrapPad Software).

## Acknowledgments

We thank RIKEN BSI-Olympus Collaboration Center for the technical assistance with the FRAP experiment. This work was supported by the JSPS KAKENHI (26650070 and 17K07399 to M. T., 16K07455 to T. O., 15K18482 and 17K15068 to T.N., 16K15095 to M.T.K., 25116009 and 16H04744 to N. S., 15H04707 to M. N., 26251003 to T. H., 15H05929 to N.I.). M.T.K. was supported by a grant from the Mochida Memorial Foundation for Medical and Pharmaceutical Research, the SGH Foundation, the Sumitomo Foundation, and the Canon Foundation.

The authors declare no competing financial interests.

Author contributions: M. Takagi designed and performed most of the experiments, generated cell lines and constructs, and analyzed data. C. Sakamoto and N. Saitoh performed the wndchrm analysis, and T. Ono and M. Nakao contributed to the wndchrm analysis. T. Natsume and M.T. Kanemaki gave advice to M. Takagi on the AID system. N. Imamoto surpervised the entire study. M. Takagi and T. Hirano wrote the paper with input from all authors.

## Supplemental materials

Fig. S1 shows the characterization of AID14 and AID15. Fig. S2 shows the FRAP analysis of Ki-67 in the presence or absence of SMC2. Fig. S3 shows the live observation of AID35 treated with IAA after the establishment of mitotic chromosome structure. Fig. S4 shows the localization of α-tubulin, Hec1 and topo IIα in cells devoid of SMC2. Fig. S5 shows the localization of centromere/kinetochore-associated proteins in cells devoid of both Ki-67 and SMC2. Table S1 lists the AID cell lines. Table S2 lists the primers used for genomic PCR. Table S3 lists antibodies used in this study.

## Supplementary figure legends

**Figure S1.**
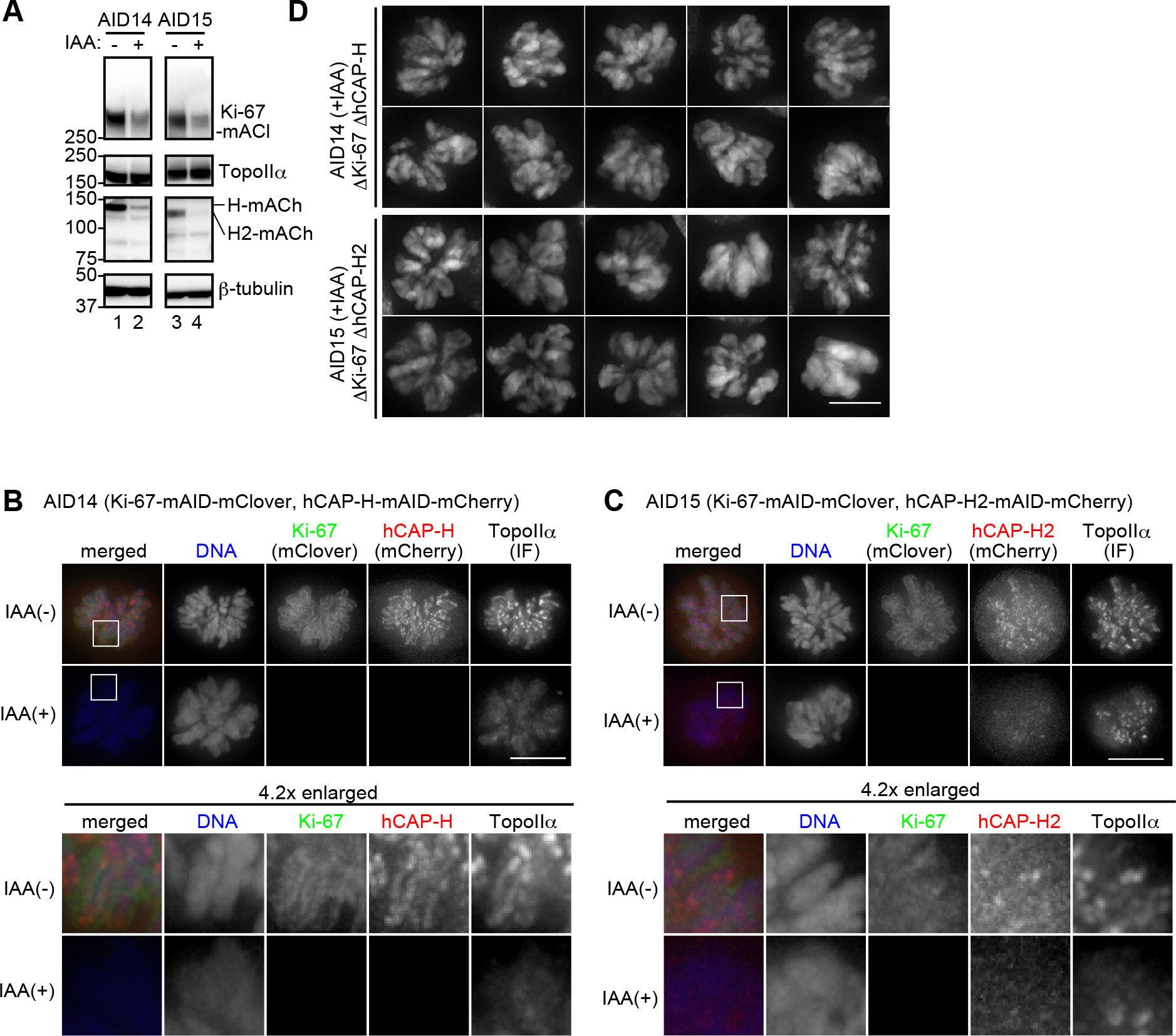
Loss of the structural integrity of mitotic chromosomes in cells devoid of Ki-67 and hCAP-H or cells devoid of Ki-67 and hCAP-H2. (A) Immunoblot analysis of AID14 and AID15 cells prepared as illustrated in Figure 1A. Membranes were probed with specific antibodies against the indicated proteins. (B-C) Immunofluorescence analysis of AID14 and AID15 cells prepared in the absence (-) or presence (+) of IAA. Ki-67 and hCAP-H/H2 were detected by the fluorescence of mClover and mCherry, respectively, fused to their C-terminal ends. Topo IIα were detected with indirect immunofluorescence (IF) using a specific antibody. DNA was counterstained with Hoechst 33342. The areas indicated by the white squares are 4.2-times enlarged and shown on the bottom. (D) Representative images of mitotic chromosomes formed in AID14 and AID15 treated with IAA. Scale bars, 10 ¼m.

**Figure S2.**
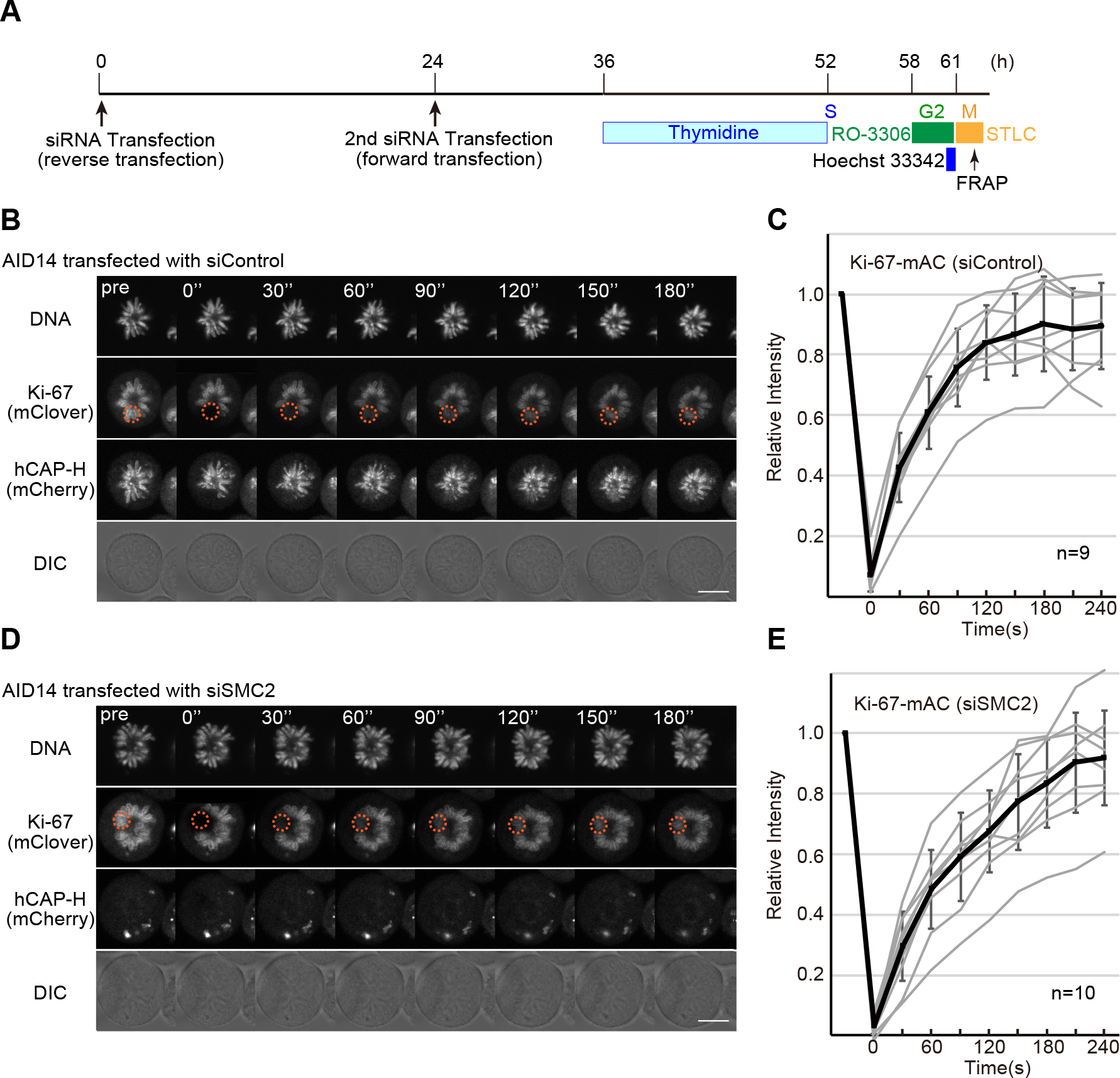
FRAP analysis of Ki-67 in the presence or absence of SMC2. (A) Schematic diagram of the cell preparation protocol. AID14 cells were transfected with control siRNA (siControl) or siRNA against SMC2 (siSMC2) twice, the first time at time 0 by a reverse transfection method and the second time at 24 h by a forward transfection method, and then processed for synchronization into mitosis. Thymidine (2 mM), RO-3306 (10 μM), Hoechst 33342 (100 ng/ml), and STLC (10 μM) were added and/or washed out at the indicated time points. (B-E) FRAP on Ki-67-mACl in AID14 transfected with siControl (B-C) or siSMC2 (D-E). (B, D) Representative image sets. The areas marked with red dashed circles were bleached. Time relative to the bleach point are indicated. (C, E) Recovery curves of Ki67-mACl fluorescence. Data from single cells were draw in grey. Thick curves and bars display mean ±SD. In AID14 transfected with siControl, Ki-67-mACl was highly mobile showing ~92% recovery in 240 s after bleaching, a result consistent with the previous observation on EGFP-Ki-67 transiently expressed in HeLa cells (Saiwaki et al., 2005). In AID14 transfected with siSMC2, depletion of SMC2 was indirectly monitored by the disappearance of hCAP-H-mACh fluorescence from mitotic chromosomes. We found that the FRAP of Ki-67-mACl in the condensin-depleted cells was indistinguishable from that observed in the control cells. Scale bars, 10 μm.

**Figure S3.**
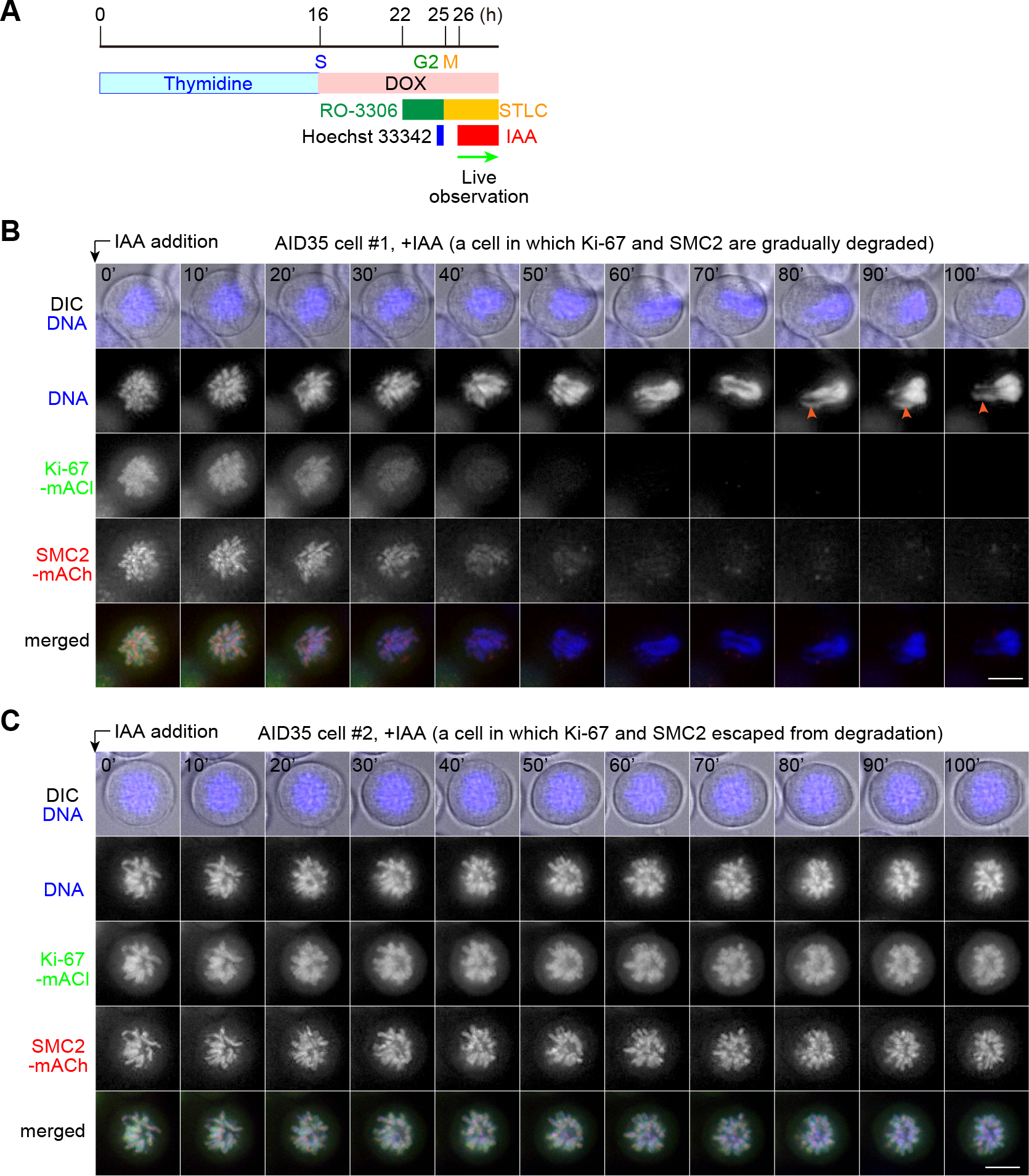
Rapid loss of the structural integrity of mitotic chromosomes in cells devoid of both Ki67 and SMC2 even after their assembly is complete. (A) Schematic diagram of the cell preparation protocol. Thymidine (2 mM), DOX (2 μg/ml), RO-3306 (10 μM), STLC (10 μM), Hoechst 33342 (100 ng/ml), and IAA (0.5 mM) were added and/or washed out at the indicated time points. AID-tagged proteins are subjected to proteasome-mediated degradation upon the treatment of cells with IAA. (B-C) AID35 cells, HCT116-based cells expressing Ki-67-mACl and SMC2-mACh, were filmed at 10-min intervals over 100 minutes. In the cell #1 (B), Ki-67-mACl and SMC2-mACh were degraded over time as intended, and mitotic chromosomes accordingly lost the structural integrity. The characteristic protrusions are marked with arrowheads. The cell #2 (C), in which both proteins escaped from degradation during the imaging period, is presented here as a control. DIC: differential interference contrast. Shown here are representative image sets out of more than eight image sets captured. Scale bars, 10 μm.

**Figure S4.**
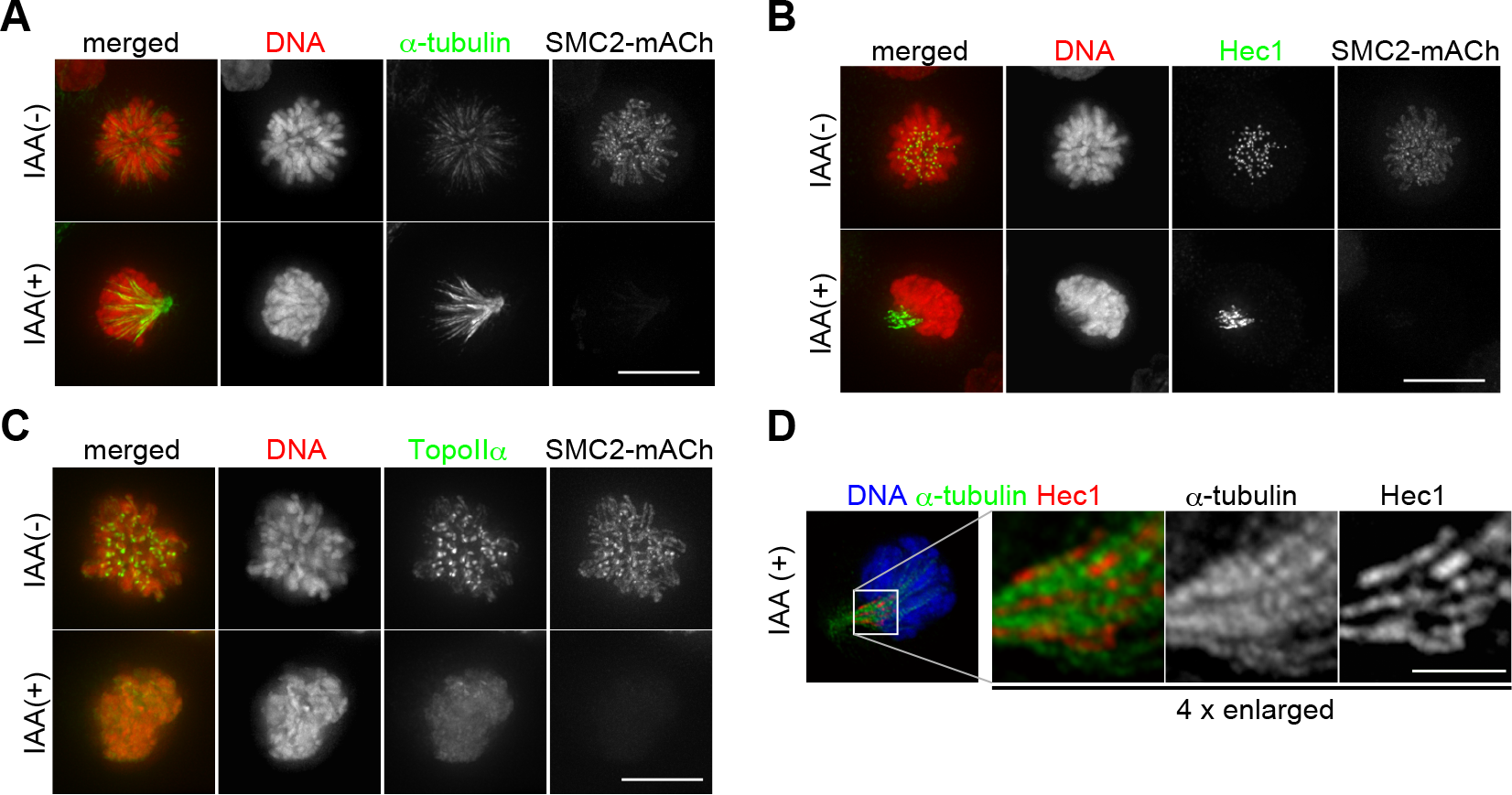
Behavior of chromosomal and non-chromosomal markers in cells devoid of SMC2. AID30 cells were prepared in the absence (-) or presence (+) of IAA according to the protocol depicted in Figure 2 A, and processed for immunofluorescence using antibodies against α-tubulin (A), Hec1 (B), topo II α (C), or α-tubulin and Hec1 (D). Ki-67 and SMC2 were detected via the fluorescence of mClover and mCherry, respectively, fused to their C-terminal ends. DNA was counterstained with Hoechst 33342. (D) The area indicated by the white square is enlarged four times and shown on the right. Note that, in the absence of SMC2, microtubules (green) were bundled and Hec1 (red) was localized along the microtubule bundles. Scale bars, 10 μm.

**Figure S5.**
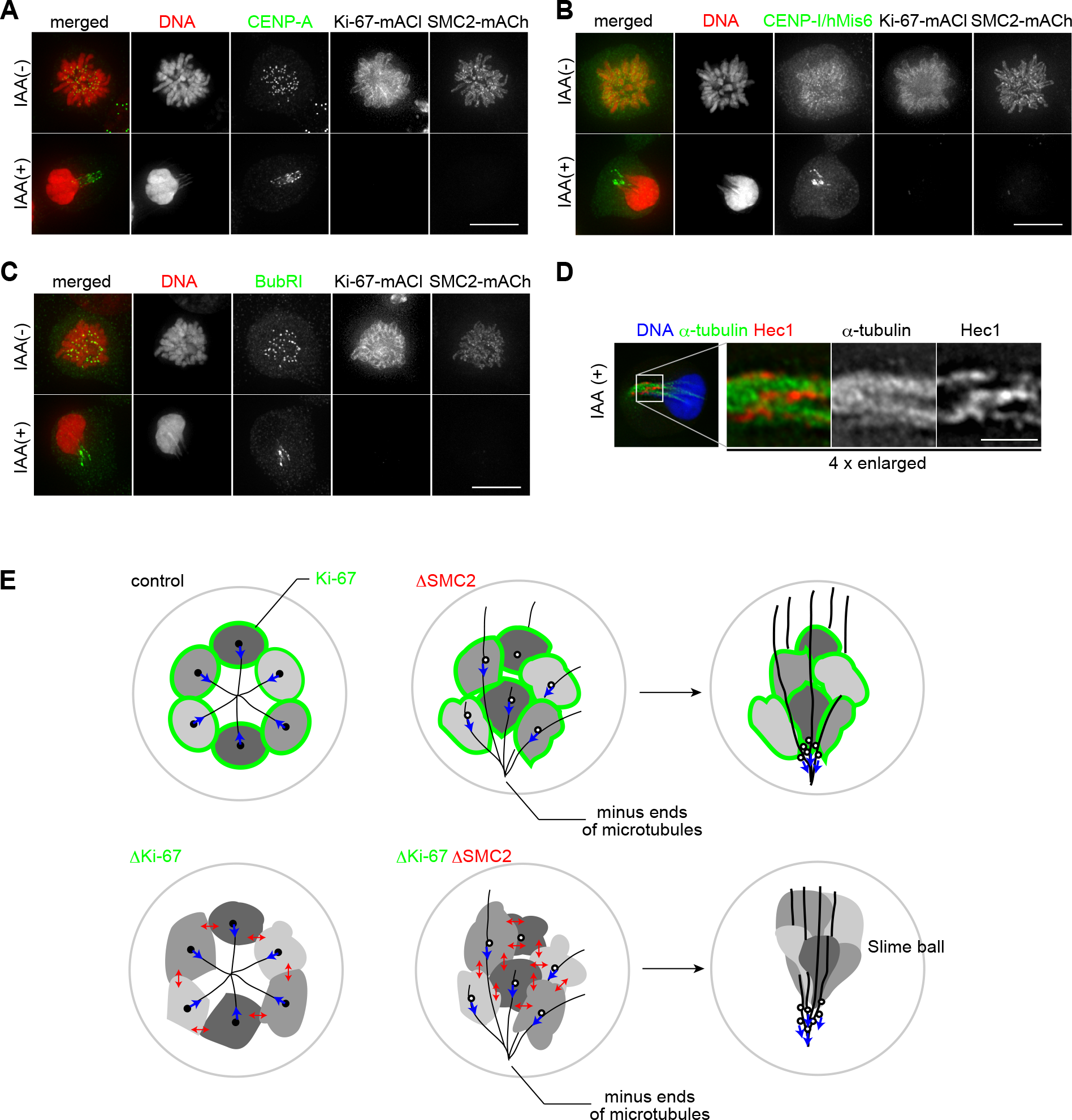
Behavior of centromere/kinetochore-associated proteins in cells devoid of both Ki-67 and SMC2. AID35 cells were treated in the absence (-) or presence (+) of IAA according to the protocol depicted in Figure 2 A, and processed for immunofluorescence using antibodies against CENP-A (A), CENP-I/hMis6 (B), BubR1 (C), or α-tubulin and Hec1 (D). Mouse anti-CENP-A monoclonal antibody (3-19, MBL) was used at 1:200 dilution in combination with goat anti-Mouse IgG (H+L), Alexa Fluor 647 (Thermo Fisher Science). Rat anti-CENP-I/hMis6 polyclonal antibody (PD032, MBL) was used at 1:100 dilution in combination with goat anti-Rat IgG (H+L), Alexa Fluor 647 (Thermo Fischer Science). Mouse anti-BubR1 monoclonal antibody (8G1, MBL) was used at 1:400 dilution in combination with goat anti-Mouse IgG (H+L), Alexa Fluor 647 (Thermo Fisher Science). Ki-67 and SMC2 were detected by the fluorescence of mClover and mCherry, respectively, fused to their C-terminal ends. DNA was counterstained with Hoechst 33342. Note that a certain amount of CENP-I/hMis6 was detected on the surface of mitotic chromosomes (B). (D) The area indicated by the white square is enlarged four times and shown on the right. Note that, in the absence of both Ki-67 and SMC2, microtubules (green) were bundled and Hec1 (red) was localized along the microtubule bundles. Scale bars, 10 [im for A-C and 2.5 im for D. (E) A model for the collapse of mitotic chromosome architecture upon degradation of both Ki-67 and SMC2 in STLC-treated cell. Paired sister mitotic chromosomes in control cells are shown in single ovoid with one kinetochore (black dot) for simplicity. The force exerted on kinetochores is depicted by the blue arrows. Upon depletion of Ki-67 (green), chromosomes come to closer and start to interfere mutually (depicted with red bidirectional arrows). Microtubules (black lines) display radial array even in the absence of Ki-67. Upon depletion of SMC2, the chromosome architecture is disturbed but not collapsed completely. At the same time, microtubules lose the symmetric array and become bundled, and the fashion of microtubule attachment to kinetochore appears to be changed from “end-on” to “lateral”. Under the condition where the effects of Ki-67-and SMC2-depletions are overlapped, the architecture of mitotic chromosome is severely collapsed. We speculate that the collapse might be accelerated by the unidirectional pulling force exerted on kinetochores along bundled microtubules. Kinetochores are denoted by two different symbols (filled or hollow circles) to express the difference in their properties. Whereas kinetochores denoted by filled circles can establish the end-on attachments of microtubules, kinetochores denoted by hollow circles tend to establish the lateral attachment to microtubules.

**Table S1.**
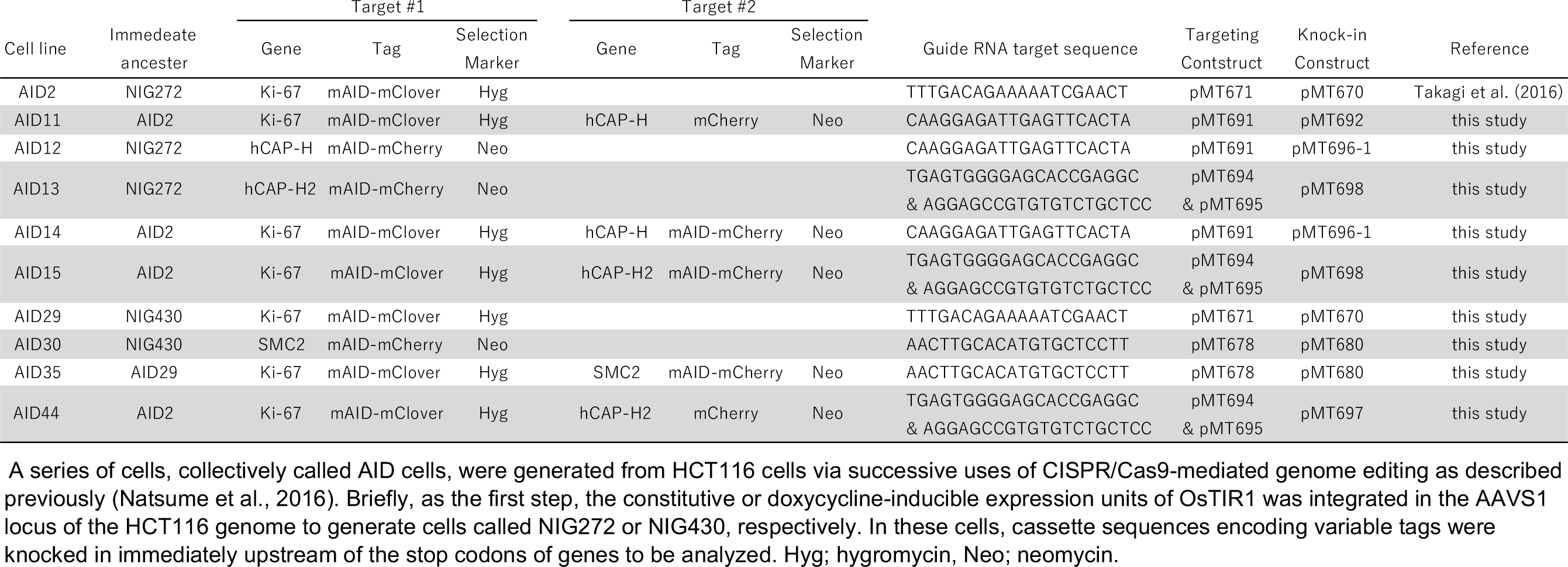
**AID cell lines used in this study.** A panel of cell lines, collectively called AID cells, were generated from HCT116 cells via successive CRISPR/Cas9-mediated genome editing as described previously (Natsume et al., 2016). Briefly, as the first step, the constitutive or doxycycline-inducible expression unit of *Os*TIR1 was integrated in the AAVS1 locus of the HCT116 genome to generate cells called NIG272 or NIG430, respectively. In these cells, cassette sequences encoding variable tags were knocked-in immediately upstream of the stop codons of genes to be analyzed. Hyg; hygromycin, Neo; neomycin.

**Table S2.**
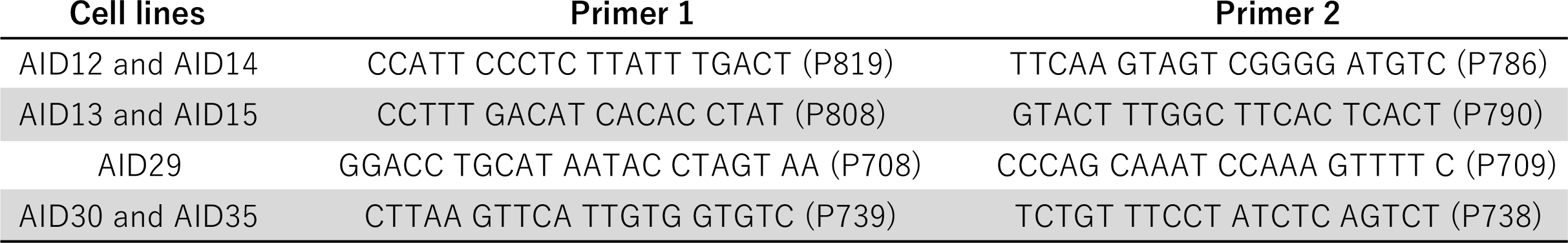
**Primers used for genomic PCR.**

**Table S3.**
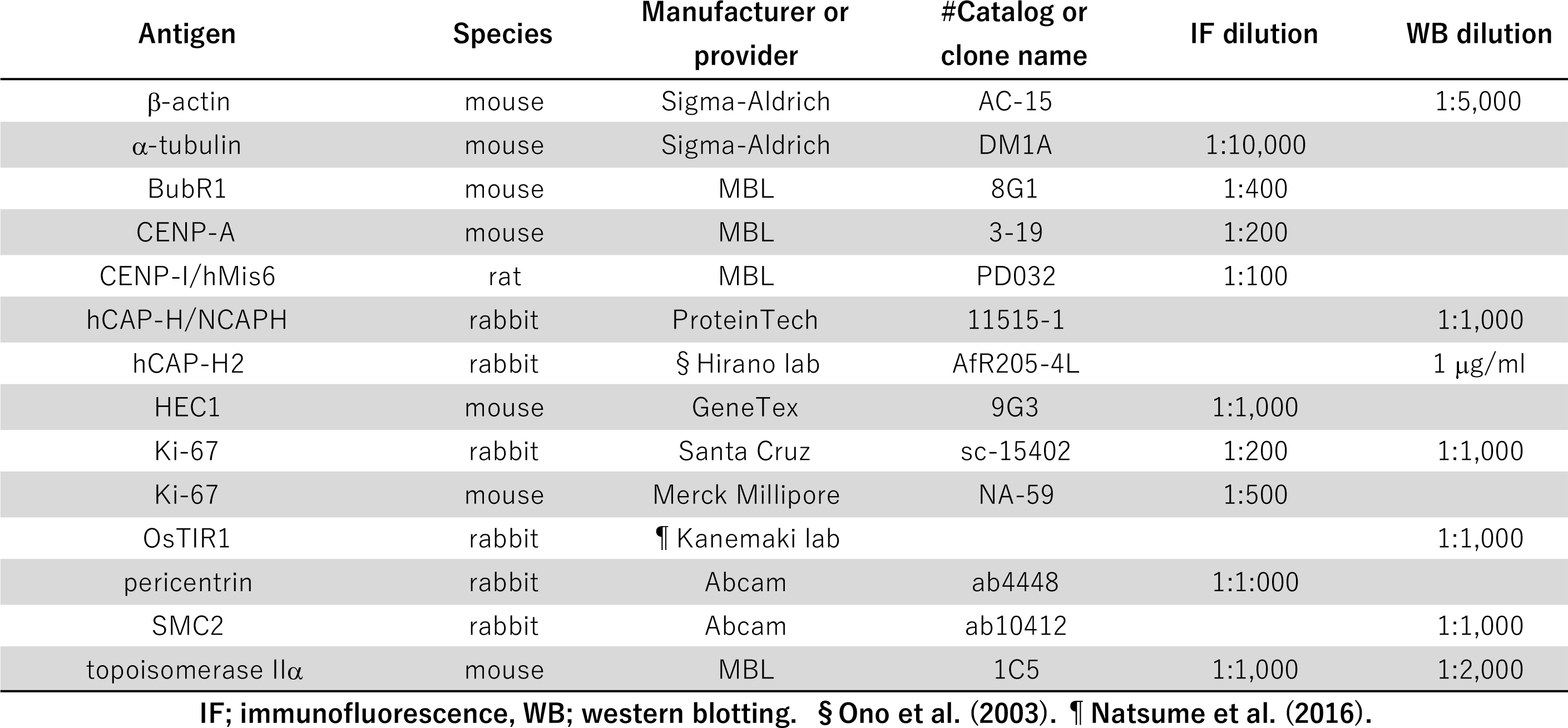
**Antibodies used in this study.**

## References

Banani, S.F., H.O. Lee, A.A. Hyman, and M.K. Rosen. 2017. Biomolecular condensates: organizers of cellular biochemistry. Nat Rev Mol Cell Biol. 18:285–298.

Booth, D.G., A.J. Beckett, O. Molina, I. Samejima, H. Masumoto, N. Kouprina, V. Larionov, I.A. Prior, and W.C. Earnshaw. 2016. 3D-CLEM Reveals that a Major Portion of Mitotic Chromosomes Is Not Chromatin. Mol Cell. 64:790–802.

Booth, D.G., M. Takagi, L. Sanchez-Pulido, E. Petfalski, G. Vargiu, K. Samejima, N. Imamoto, C.P. Ponting, D. Tollervey, W.C. Earnshaw, and P. Vagnarelli. 2014. Ki-67 is a PP1-interacting protein that organises the mitotic chromosome periphery. eLife. 3:e01641.

Cuylen, S., C. Blaukopf, A.Z. Politi, T. Muller-Reichert, B. Neumann, I. Poser, J. Ellenberg, A.A. Hyman, and D.W. Gerlich. 2016. Ki-67 acts as a biological surfactant to disperse mitotic chromosomes. Nature. 535:308–312.

Gassmann, R., P. Vagnarelli, D. Hudson, and W.C. Earnshaw. 2004. Mitotic chromosome formation and the condensin paradox. Exp Cell Res. 296:35–42.

Gerlich, D., T. Hirota, B. Koch, J.M. Peters, and J. Ellenberg. 2006. Condensin I stabilizes chromosomes mechanically through a dynamic interaction in live cells. Current biology: CB. 16:333–344.

Green, L.C., P. Kalitsis, T.M. Chang, M. Cipetic, J.H. Kim, O. Marshall, L. Turnbull, C.B. Whitchurch, P. Vagnarelli, K. Samejima, W.C. Earnshaw, K.H. Choo, and D. F. Hudson. 2012. Contrasting roles of condensin I and condensin II in mitotic chromosome formation. Journal of cell science. 125:1591–1604.

Hirano, T. 2016. Condensin-Based Chromosome Organization from Bacteria to Vertebrates. Cell. 164:847–857.

Hirota, T., D. Gerlich, B. Koch, J. Ellenberg, and J.M. Peters. 2004. Distinct functions of condensin I and II in mitotic chromosome assembly. Journal of cell science. 117:6435–6445.

Johnston, J., W.B. Iser, D.K. Chow, I.G. Goldberg, and C.A. Wolkow. 2008. Quantitative image analysis reveals distinct structural transitions during aging in Caenorhabditis elegans tissues. PLoS One. 3:e2821.

Maresca, T.J., B.S. Freedman, and R. Heald. 2005. Histone H1 is essential for mitotic chromosome architecture and segregation in Xenopus laevis egg extracts. J Cell Biol. 169:859–869.

Mazumdar, M., J.H. Lee, K. Sengupta, T. Ried, S. Rane, and T. Misteli. 2006. Tumor formation via loss of a molecular motor protein. Current biology: CB. 16:1559–1564.

Natsume, T., T. Kiyomitsu, Y. Saga, and M.T. Kanemaki. 2016. Rapid Protein Depletion in Human Cells by Auxin-Inducible Degron Tagging with Short Homology Donors. Cell Rep. 15:210–218.

Ono, T., Y. Fang, D.L. Spector, and T. Hirano. 2004. Spatial and temporal regulation of Condensins I and II in mitotic chromosome assembly in human cells. Mol Biol Cell. 15:3296–3308.

Ono, T., C. Sakamoto, M. Nakao, N. Saitoh, and T. Hirano. 2017. Condensin II plays an essential role in reversible assembly of mitotic chromosomes in situ. Mol Biol Cell.

Orlov, N., L. Shamir, T. Macura, J. Johnston, D.M. Eckley, and I.G. Goldberg. 2008. WND-CHARM: Multi-purpose image classification using compound image transforms. Pattern Recognit Lett. 29:1684–1693.

Saiwaki, T., I. Kotera, M. Sasaki, M. Takagi, and Y. Yoneda. 2005. In vivo dynamics and kinetics of pKi-67: transition from a mobile to an immobile form at the onset of anaphase. Exp Cell Res. 308:123–134.

Samejima, K., I. Samejima, P. Vagnarelli, H. Ogawa, G. Vargiu, D.A. Kelly, F. de Lima Alves, A. Kerr, L.C. Green, D.F. Hudson, S. Ohta, C.A. Cooke, C.J. Farr, J. Rappsilber, and W.C. Earnshaw. 2012. Mitotic chromosomes are compacted laterally by KIF4 and condensin and axially by topoisomerase IIalpha. J Cell Biol. 199:755–770.

Scholzen, T., and J. Gerdes. 2000. The Ki-67 protein: from the known and the unknown. J Cell Physiol. 182:311–322.

Shintomi, K., T.S. Takahashi, and T. Hirano. 2015. Reconstitution of mitotic chromatids with a minimum set of purified factors. Nat Cell Biol. 17:1014–1023.

Silk, A.D., A.J. Holland, and D.W. Cleveland. 2009. Requirements for NuMA in maintenance and establishment of mammalian spindle poles. J Cell Biol. 184:677–690.

Takagi, M., T. Natsume, M.T. Kanemaki, and N. Imamoto. 2016. Perichromosomal protein Ki67 supports mitotic chromosome architecture. Genes Cells. 21:1113–1124.

Takagi, M., Y. Nishiyama, A. Taguchi, and N. Imamoto. 2014. Ki67 antigen contributes to the timely accumulation of protein phosphatase 1gamma on anaphase chromosomes. The Journal of biological chemistry. 289:22877–22887.

Takahashi, M., T. Wakai, and T. Hirota. 2016. Condensin I-mediated mitotic chromosome assembly requires association with chromokinesin KIF4A. Genes Dev. 30:1931–1936.

Tokunaga, K., N. Saitoh, I.G. Goldberg, C. Sakamoto, Y. Yasuda, Y. Yoshida, S. Yamanaka, and M. Nakao. 2014. Computational image analysis of colony and nuclear morphology to evaluate human induced pluripotent stem cells. Sci Rep. 4:6996.

Uhlmann, F. 2016. SMC complexes: from DNA to chromosomes. Nat Rev Mol Cell Biol. 17:399–412.

Vanneste, D., M. Takagi, N. Imamoto, and I. Vernos. 2009. The role of Hklp2 in the stabilization and maintenance of spindle bipolarity. Current biology: CB. 19:1712–1717.

